# TREX reveals proteins that bind to specific RNA regions in living cells

**DOI:** 10.1101/2023.06.30.547259

**Authors:** Martin Dodel, Giulia Guiducci, Maria Dermit, Sneha Krishnamurthy, Lovorka Stojic, Faraz K. Mardakheh

## Abstract

Different regions of RNA molecules can often engage in specific interactions with distinct RNA-binding proteins (RBPs), giving rise to diverse modalities of RNA regulation and function. However, there are currently no methods for unbiased identification of RBPs that interact with specific RNA regions in living cells under endogenous settings. Here, we introduce TREX (Targeted RNase H-mediated extraction of crosslinked RBPs), a highly sensitive approach for identifying proteins that directly bind to specific RNA regions in living cells. We demonstrate that TREX outperforms existing methods in identifying known interactors of *U1* snRNA, and reveals endogenous region-specific interactors of *NORAD* lncRNA. Using TREX, we generated a comprehensive region-by-region interactome for *45S* rRNA, uncovering both established and novel interactions that regulate ribosome biogenesis. With its applicability to any RNA in any cell-type, TREX is the first RNA-centric tool for unbiased positional mapping of endogenous RNA-protein interactions in living cells.

## Introduction

The fate and function of every RNA molecule in living organisms is defined by its interactions with RNA binding proteins (RBPs). RNA-RBP interactions, therefore, play a fundamental role in regulating all aspects of cell behavior and homeostasis. Several methods have been developed for deciphering specific RNA-RBP interactions in living cells. These can be broadly divided into protein-centric approaches, which reveal the RNAs that are bound by a specific RBP, or RNA-centric approaches, which reveal the RBPs that bind to a specific RNA (Hafner et al., 2021). Crosslinking and Immunoprecipitation (CLIP)-based methods are amongst the most powerful and widely used protein-centric approaches, which leverage the sensitivity of next-generation sequencing to not only profile the compendium of RNAs that directly interact with a given RBP, but also enable the mapping of the precise binding sites of the RBP on its target RNAs (Hafner et al., 2010; Konig et al., 2010; Ule et al., 2003; Van Nostrand et al., 2016). Transcriptome-wide binding sites of hundreds of RBPs have been profiled using different CLIP-based approaches (Van Nostrand et al., 2020). A more recent protein-centric approach for RNA-RBP profiling involves the fusion of RBPs with RNA-editing enzymes, followed by transcriptome-wide identification of modified RNAs via next-generation sequencing (Brannan et al., 2021; McMahon et al., 2016). By eliminating the need for immunoprecipitation, these methods have pushed the limits of protein-centric RNA-RBP profiling down to the single-cell level (Brannan *et al*., 2021).

In contrast, RNA-centric profiling of RNA-RBP interactions has remained more challenging. The most common methods use crosslinking followed by RNA affinity capture and mass spectrometry (MS) analysis, to purify and identify proteins that directly interact with an RNA target (Hafner *et al*., 2021). For example, biotinylated antisense oligonucleotides coupled with streptavidin conjugated beads have been utilized to profile the interactomes of endogenous *U1*, *XIST*, *NORAD*, *18S* and *RMRP* non-coding RNAs, amongst others (Chu et al., 2015; Desideri et al., 2020; McHugh et al., 2015; Munschauer et al., 2018). Low efficiency has been a major caveat of these pulldown-based methods, often requiring large numbers of cells per experiment for successful identification of interacting proteins (McHugh and Guttman, 2018). An alternative strategy involves the use of proximity-based labeling enzymes, such as Biotin Ligase BirA (BioID) or Ascorbate Peroxidase (APEX), which can be targeted to a specific RNA of interest (Han et al., 2020; Mukherjee et al., 2019; Ramanathan et al., 2018; Tsue et al., 2023; Zhang et al., 2020). Subsequently, the locally labeled proteins are captured and identified by MS analysis. A notable limitation of this approach is the inclusion of indirectly associated proteins that are in close physical proximity to the target RNA. Most importantly, both RNA affinity capture and proximity-based labeling methods fail to provide any insight into the specific regions of the target RNA that are responsible for mediating interactions with individual proteins.

To address these shortcomings, we developed TREX. Short for ‘Targeted RNase H-mediated Extraction of crosslinked(X) RBPs’, TREX provides a highly efficient method to extract RBPs that are bound to a specific RNA sequence, followed by their identification by quantitative MS. Unlike previous RNA-centric methods, TREX can map endogenous RNA-protein interactions in a region-specific manner. We first benchmarked TREX against the full length of *U1* small nuclear RNA (snRNA), the well-characterized RNA component of the U1 spliceosome complex (Pomeranz Krummel et al., 2009), demonstrating that it outperforms existing antisense oligo-based pulldown methods for revealing known *U1* interactions. Next, we applied TREX to the most conserved segment of *NORAD* long noncoding RNA (lncRNA), highlighting its versatility in probing region-specific interactions in endogenous settings. Finally, we used TREX to generate a detailed region-by-region interactome for *45S* pre-ribosomal RNA (rRNA) gene products. This mapping unveiled many known as well as several previously unknown interactors, providing insights into the intricate regulation of ribosome biogenesis and composition that is organized by distinct sections of *45S* pre-rRNA. Our findings establish TREX as a robust and versatile RNA-centric approach for revealing region-specific RNA-RBP interactions. We envisage that TREX will advance our understanding of diverse post-transcriptional regulatory processes in many cell types and organisms.

## Results

### Combining organic phase separation with targeted RNase H-mediated degradation

Several recent studies have reported that UV-C crosslinking in combination with Acid Guanidinium Thiocyanate-Phenol-Chloroform extraction can be used to isolate and purify RBPs that directly bind RNA (Queiroz et al., 2019; Trendel et al., 2019; Urdaneta et al., 2019). In this approach, UV-C treatment is employed to covalently crosslink RNA molecules to their interacting RBPs in living cells. Subsequent cell lysis in acidic guanidinium thiocyanate-phenol mixture (commonly referred to as TRIZOL) results in the disruption of lipid membranes and denaturation of macromolecules. The resulting lysate is then subjected to organic phase separation by addition of chloroform, leading to the partitioning of the free RNA molecules to the upper aqueous phase and the free proteins to the lower organic phase. Covalently linked RNA-RBP complexes, however, cannot partition into either phase, but instead form an insoluble interface that can be isolated and subjected to shotgun proteomics analysis for comprehensive profiling of the RNA-bound proteome (Queiroz *et al*., 2019; Trendel *et al*., 2019).

In TREX, we utilize a similar approach to isolate RNA-RBP complexes, but the interface is subjected to annealing of tiling antisense DNA oligonucleotides that are fully complementary to an RNA sequence of interest. The targeted RNA sequence, now hybridized to DNA, is specifically degraded by the addition of RNase H, an enzyme that specifically degrades RNA molecules that are hybridized to DNA, resulting in the release of any crosslinked RBPs. Subsequently, a second organic phase separation is used to partition the released RBPs into the organic phase, allowing their extraction from the remaining RNA-bound RBPs that stay in the interface. The released RBPs are then identified by quantitative MS analysis (Fig. 1A). We confirmed that intact total RNA from the interface can be effectively solubilized for downstream processing (Fig. S1A and S1B). Moreover, upon RNase H addition, tiling antisense oligonucleotides against several different types of RNA targets were confirmed to trigger the specific depletion of their corresponding targets in a dose dependent manner (Fig. S1C-F). These findings indicate that the crucial steps of TREX, including interface solubilization and subsequent RNase H-mediated target degradation, operate efficiently to recover and selectively deplete protein-bound RNA sequences of interest.

**Figure 1:**
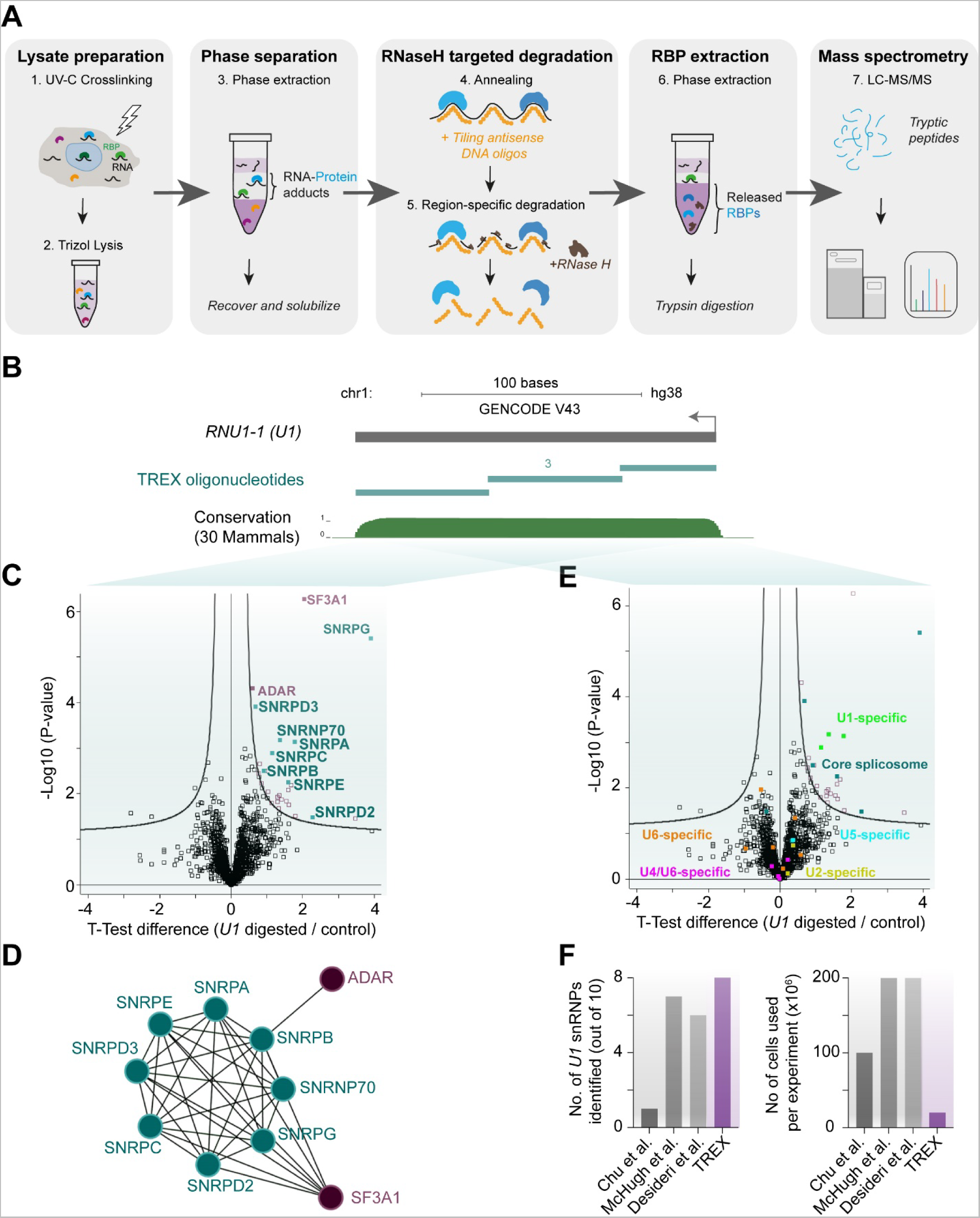
TREX reveals proteins that bind to specific RNA sequences in living cells. **(A)** Experimental scheme of TREX. **(B)** Schematic representation of *U1* snRNA primary sequence (annotated as *RNU1-1* in RefSeq) with the associated UCSC Genome Browser tracks for mammalian conservation (PhastCons). The tiling antisense DNA oligonucleotides used for depletion of *U1* in TREX (total: 3) are depicted on the tracks. Scale = 100 bases. **(C)** Volcano plot of the two-sample t-test comparison of *U1* digested vs undigested TREX samples, showing significant enrichment of the U1 SNRNP complex members, along with SF3A1 and ADAR in the digested samples. Curved lines mark the significance boundary (FDR = 0.05, S0 = 0.1). **(D)** Interaction network analysis of *U1*-bound proteins identified by TREX, using the STRING physical interactions database (Szklarczyk et al., 2015). **(E)** Volcano plot of the two-sample t-test comparison of *U1* digested vs undigested TREX samples, with marking the core spliceosome as well as each identified specific U SNRNP complex protein members. Only *U1*-specific proteins, as well as the core spliceosome components, are found as significantly enriched in the *U1* digested TREX samples. Curved lines mark the significance boundary (FDR = 0.05, S0 = 0.1). **(F)** Comparison of the number of cells used per experiment, against the number of known U1 SNRNP proteins identified, in TREX versus three previous *U1* RNA affinity capture-MS studies (Chu *et al*., 2015; Desideri *et al*., 2020; McHugh *et al*., 2015).

### TREX outperforms existing RNA pulldown-based approaches in revealing the direct interactions of *U1* snRNA

Next, we benchmarked TREX by applying it to analyze the direct binding partners of *U1* snRNA (Fig. 1B), the RNA component of the *U1* small nuclear ribonucleoprotein particle (snRNP) (Pomeranz Krummel *et al*., 2009), whose protein interactions are well-characterized. Human *U1* snRNP contains 10 proteins, seven of which are core spliceosome components (SNRPB, SNRPD1, SNRPD2, SNRPD3, SNRPE, SNRPF, and SNRPG), while the remaining three (SNRPA, SNRPC, and SNRNP70) are *U1*-specific factors. For each biological replicate, 10 million HCT116 human colorectal carcinoma cells were subjected to TREX analysis. As a negative control, a parallel preparation was performed for each replicate, but without the addition of RNase H. To assess the efficiency of the RNase H-mediated degradation, RNA was isolated from a portion of each TREX reaction before the second organic phase separation (step five - Fig. 1A). The levels of *U1* RNA were assessed by RT-qPCR, confirming its effectively complete removal in the RNase H-treated samples (Fig. S1G). The specificity of RNase H-mediated degradation was also evaluated through whole-transcriptome RNA-sequencing analysis of the same RNA samples, which revealed no other transcripts were significantly impacted (Fig. S1H). The remainder of each reaction was subjected to the second organic phase separation to isolate the released proteins, which were subsequently recovered, trypsin digested, and analyzed by quantitative MS (Steps 6 and 7 - Fig. 1A). Principal component analysis (PCA) of the results indicated high reproducibility amongst the RNase H-treated biological replicates (Fig. S1I). Overall, 27 proteins were identified as significant direct interactors of *U1* (Fig. 1C and Dataset S1). Amongst these, eight out of the 10 U1 snRNP components, along with SF3A1 and ADAR, two other known interactors of U1 snRNP (Agranat et al., 2008; Sharma et al., 2014), were identified (Fig. 1C, 1D, and Dataset S1). Importantly, proteins specific to other snRNP complexes such as *U2, U4, U5,* and *U6* snRNPs were not amongst the significant binding proteins, showcasing the highly specific nature of TREX (Fig. 1E and Dataset S1).

Since the *U1* interactome has been previously investigated in a number of RNA pulldown-based studies, we were able to evaluate the performance of TREX against these studies (Chu *et al*., 2015; Desideri *et al*., 2020; McHugh *et al*., 2015). While all previous pulldown-based analyses of *U1* used 100 million or more cells per replicate experiment, only 10 million cells per replicate were required for TREX (Fig. 1F). Nonetheless, TREX could identify eight out of the 10 *U1* snRNP proteins, which is more than all the other pulldown-based studies (Fig. 1F). Together, these results demonstrate that from a fraction of the input material, TREX is capable of achieving superior results in terms of revealing known interactions.

### TREX identifies the direct interactors of the ND4 segment of *NORAD* lncRNA

Encouraged with the TREX analysis of full-length *U1* snRNA, we next set out to investigate the capacity of TREX to decipher region-specific interactions. To address this, we focused on *NORAD* (noncoding RNA activated by DNA damage), a 5.3 kb-long lncRNA that is highly conserved and robustly expressed in human tissues and cell lines (Lee et al., 2016; Tichon et al., 2016). *NORAD* is known to be upregulated in response to DNA damage, and its main reported function is to safeguard genome stability. Indeed, *NORAD* depletion in HCT116 cells leads to chromosome segregation errors (Lee *et al*., 2016; Munschauer *et al*., 2018). Previous studies have indicated that *NORAD* localizes predominantly in the cytoplasm, but is also present in the nucleus (Lee *et al*., 2016; Matheny et al., 2021; Munschauer *et al*., 2018; Tichon *et al*., 2016). *NORAD* contains approximately 20 Pumilio Recognition Elements (PREs), which are binding sites for translational regulators Pumilio 1 and Pumilio 2 (PUM1 and PUM2), representing the primary known functional interactors of *NORAD (Elguindy et al., 2019; Kopp et al., 2019; Lee et al., 2016; Tichon et al., 2016)*. *NORAD* has a unique primary sequence structure consisting of five repetitive ~400 nucleotide (nt) domains known as *NORAD* Domains (ND1-ND5) (Lee *et al*., 2016). The ND4 domain represents the most conserved of the five repeated segments (Fig. 2A). Importantly, although multiple studies have assessed *NORAD* interacting proteome through different methodologies (Elguindy *et al*., 2019; Lee *et al*., 2016; Munschauer *et al*., 2018; Spiniello et al., 2018; Tichon *et al*., 2016), identification of the direct interactors of the conserved ND4 segment under endogenous conditions in living cells has not been possible up to now.

**Figure 2:**
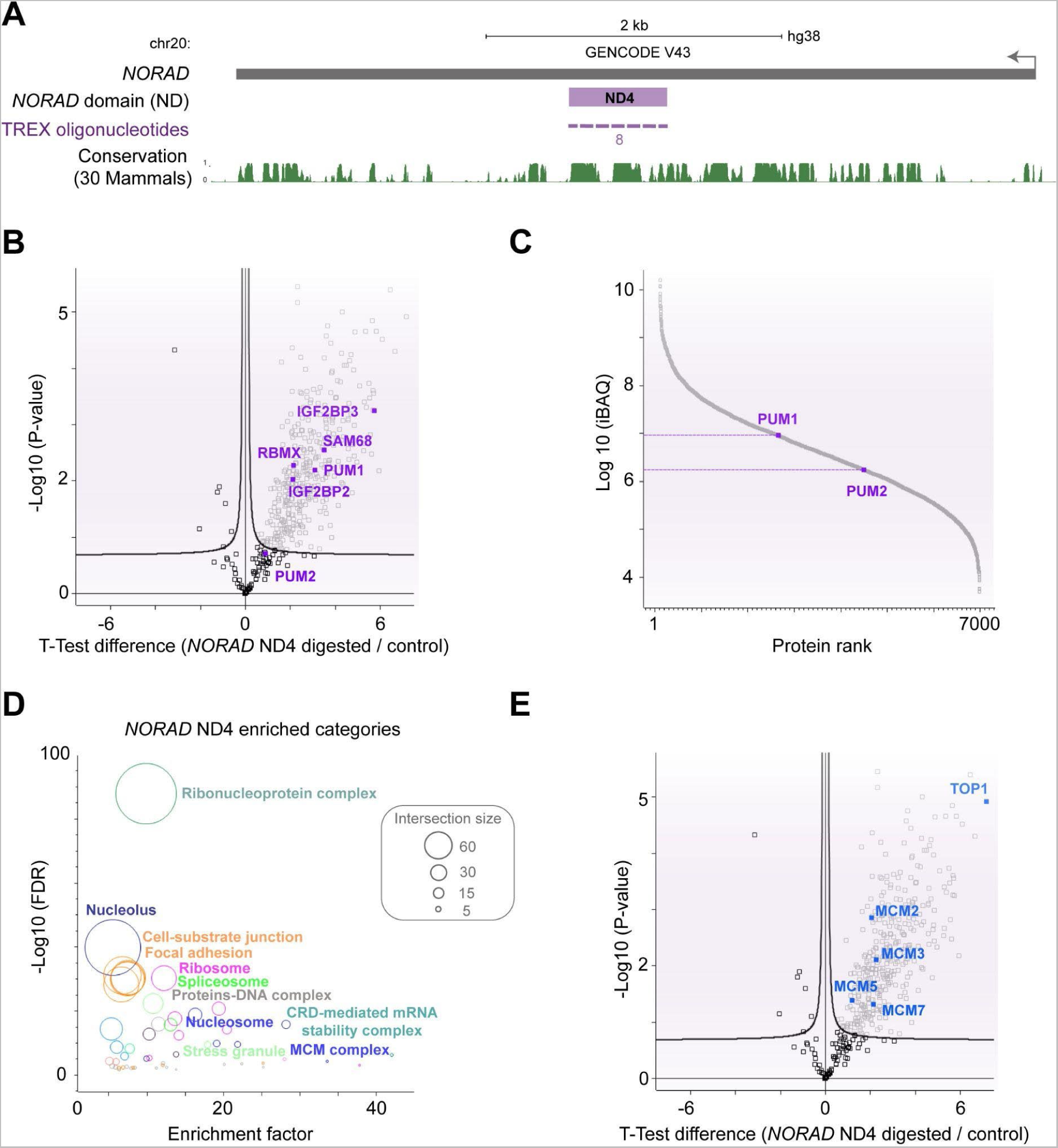
TREX defines the proteins that bind to the ND4 segment of *NORAD* lncRNA. **(A)** Schematic representation of *NORAD* lncRNA primary sequence and its ND4 segment, with the associated UCSC Genome Browser tracks for mammalian conservation (PhastCons). The tiling antisense DNA oligonucleotides used for depletion in TREX (total: 8) are depicted on the tracks. Scale = 2kilobases. **(B)** Volcano plot of the two-sample t-test comparison of *NORAD* ND4 digested vs undigested TREX samples, showing the enrichment of several known RBP interactors of *NORAD* (purple) in the digested samples. Curved lines mark the significance boundary (FDR = 0.05, S0 = 0.1). While PUM1 is a highly significant interactor, PUM2 falls just below significant cut-off. **(C)** The ranked plot of the iBAQ (Schwanhausser et al., 2011) absolute protein abundance measurements from the total proteome of HCT116, revealing PUM1 to be > 5 fold more expressed than PUM2. **(D)** Fisher’s exact test analysis of known protein categories that are over-represented amongst the *NORAD* interacting proteins (Benjamini-Hochberg FDR < 0.05). Each circle represents an enriched category from the Gene Ontology Cellular Compartments (GOCC) database, with circle size representing the number of shared proteins. **(E)** Volcano plot of the two-sample t-test comparison of *NORAD* ND4 digested vs undigested TREX samples, with TOP1, and members of the MCM helicase complex highlighted. Curved lines mark the significance boundary (FDR = 0.05, S0 = 0.1).

Thus, we employed TREX to investigate the direct interactors of *NORAD* ND4 domain. First, we validated the efficiency and specificity of the RNase H-mediated targeting by RT-qPCR, demonstrating that the ND4 region is specifically depleted without degrading a separate region of *NORAD* (Fig. S2A and S2B). Subsequently, we conducted quantitative MS analysis from 100 million HCT116 cells per replicate experiment. PCA confirmed the reproducibility amongst biological replicate samples, with the exception of one replicate from the RNase H treated samples, which was subsequently removed from further analysis (Fig. S2C). Overall, our analysis identified 360 proteins as significant direct interacting partners of the *NORAD* ND4 region. Notably, this dataset included proteins such as PUM1, SAM68, IGF2BP2/3, PABPN1 and RBMX (Fig. 2B and Dataset S2), all of which were previously reported as key *NORAD* interactors (Lee *et al*., 2016; Munschauer *et al*., 2018; Tichon *et al*., 2016; Tichon et al., 2018). PUM2 did not reach significance in our analysis (Fig. 2B and Dataset S2), possibly due to its fivefold lower expression in HCT116 cells compared to PUM1 (Fig. 2C). Gene Ontology analysis revealed diverse protein categories belonging to various nuclear as well as cytoplasmic RNA and DNA-associated proteins families (Fig. 2D). Indeed, the ND4 protein interactors seemed to be evenly split between cytoplasmic and nuclear protein families (Fig. S2D), aligning with the reported localization *NORAD* to both compartments. We also compared the overlap between our ND4 TREX data with results of other studies that have assessed the *NORAD* interacting proteome through different methods such as *in vitro* RNA pulldown coupled with MS (IVT-MS) (Lee *et al*., 2016; Tichon *et al*., 2016), hybridization-proximity coupled with MS (HyPr-MS) (Spiniello *et al*., 2018), and RNA antisense purification coupled with MS (RAP-MS) (Munschauer *et al*., 2018). We observed a highly significant overlap between our TREX results and all these studies, with ~18 to 44% of hits in these reports matching our TREX identified interactors (Fig. S2E)

Crucially, amongst novel hits, we identified topoisomerase I (TOP1) as one of the most enriched interactors of *NORAD* ND4 (Fig. 2E and Dataset S2). This finding is strongly in line with a recently described role for *NORAD* in regulating assembly of the RBMX-dependent ribonucleoprotein complex, which contains TOP1 (Munschauer *et al*., 2018). Additionally, several subunits of the MCM2-7 replication helicase, which plays a central role in DNA replication initiation and elongation, were found to interact with *NORAD* (Fig. 2D, 2E and Dataset S2). Considering that TOP1 is known to bind to active replication origins and co-purifies with the MCM complex (Gambus et al., 2006), we speculate that *NORAD*’s function in genome stability is linked to its direct binding to the TOP1-MCM complex via ND4. This finding is consistent with the observed DNA replication defects triggered by the reduction in replication fork velocity in *NORAD*-depleted cells (MM paper), resembling the phenotype associated with TOP1 loss-of-function (Tuduri et al., 2009). Collectively, our study reveals key previously unknown direct interactions between *NORAD* and the TOP1-MCM complex, demonstrating the power of TREX for robustly deciphering region-specific functional interactors of a given RNA region in endogenous settings.

### TREX reveals the collection of proteins that bind to human *45S* rRNA gene products

Next, we employed TREX to identify proteins that interact with both premature and fully processed rRNA molecules that derive from the *45S* rRNA gene in human cells. The *45S* rRNA is transcribed in the nucleolus as a >13kb long transcript containing the *18S*, *5.8S* and *28S* rRNA sequences, interspersed with four spacer sequences known as *5’ External Transcribed Spacer* (*5’ETS*), *Internal Transcribed Spacer 1* (*ITS1*), *Internal Transcribed Spacer 2* (*ITS2*), and *3’ External Transcribed Spacer* (*3’ETS*) (Fig. 3A). Through a complex series of molecular events, *45S* pre-rRNA undergoes extensive processing, resulting in the removal of spacer regions and the formation of mature ribosomal subunits containing the *18S, 5.8S*, and *28S* rRNA molecules (Dorner et al., 2023; Klinge and Woolford, 2019). To identify the complete set of proteins interacting with the RNA products of the *45S* gene, we employed TREX with tiling antisense DNA oligonucleotides spanning the entire length of *45S* pre-rRNA (Fig. 3A). As before, the efficiency of RNase H-mediated depletion of *45S* pre-rRNA was validated by RT-qPCR (Fig. S3A), while the specificity of RNase H-mediated degradation was assessed through whole-transcriptome RNA-sequencing as before, which revealed no other transcripts were significantly affected (Fig. S3B). Subsequently, we conducted quantitative MS analysis from 10 million HCT116 cells per replicate experiment. PCA of the MS data demonstrated good reproducibility among the biological replicates (Fig. S3C). We identified 160 proteins as significant direct interactors of *45S* rRNA, 97 of which were already known to interact with rRNA/ribosomes (Fig. 3B and Dataset S3). Among them were large ribosomal subunit proteins (RPLs), small ribosomal subunit proteins (RPSs), ribosome associated factors (RAFs), and known ribosome biogenesis factors (RBFs) (Fig. 3B and 3C). To further investigate the function of the remaining 63 novel interactors, we compared our dataset with hits from three RNAi screens targeting functional regulators of human ribosome biogenesis, including two genome-wide screens and one nucleolar proteome-specific screen (Badertscher et al., 2015; Dorner et al., 2022; Tafforeau et al., 2013). This analysis revealed a further nine proteins that had a role in ribosome biogenesis, despite not being known to interact with rRNA (Fig. 3B and 3C). Furthermore, protein interaction network analysis using the STRING database (Szklarczyk *et al*., 2015) showed that 153 of the 160 proteins identified can physically associate with each other (Fig. 3D). These interactions include the majority of the remaining proteins that lacked any known associations or functional links to ribosome biogenesis, suggesting that they are likely novel factors involved in rRNA-related processes. Together, these results demonstrate the ability of TREX to reveal the full repertoire of proteins that interact with *45S* rRNA gene products in living human cells.

**Figure 3:**
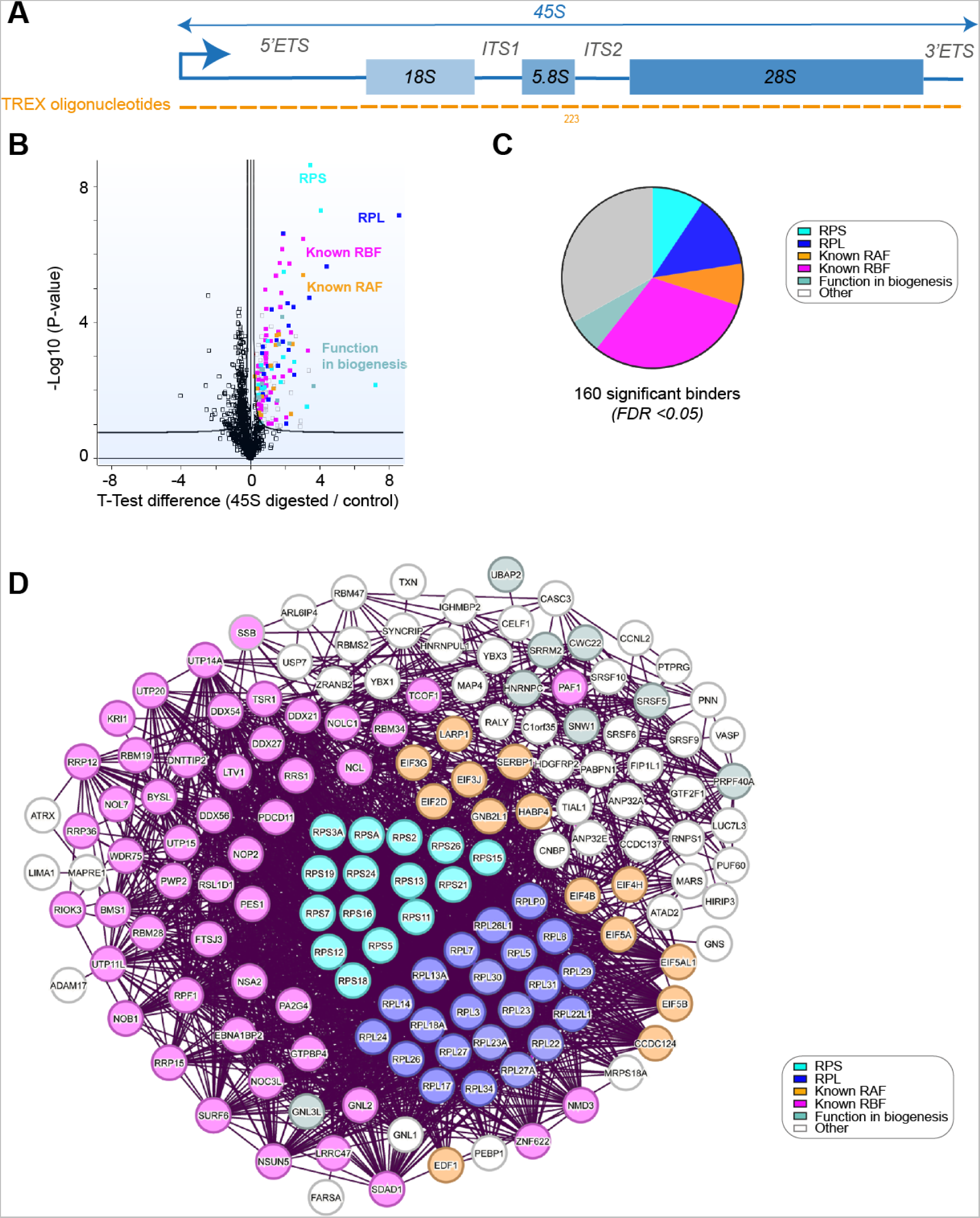
TREX captures the compendium of proteins that bind to the *45S* rRNA. **(A)** Schematic representation of *45S* rRNA primary sequence and its various segments. The tiling antisense DNA oligonucleotides used for depletion in TREX (total: 223) are depicted on the graph. **(B)** Volcano plot of the two-sample t-test comparison of *45S* rRNA digested vs undigested TREX samples, showing the enrichment of RPS (cyan) and RPL (blue) proteins, known RBFs (purple) (Dorner *et al*., 2023), known RAFs (orange), as well as previously unknown interactors suspected of involvement in human ribosome biogenesis based on the impact of their depletion (teal) (Badertscher *et al*., 2015; Dorner *et al*., 2022; Tafforeau *et al*., 2013). Curved lines mark the significance boundary (FDR = 0.05, S0 = 0.1). **(C)** Pie diagram of the composition of proteins identified in the *45S* rRNA TREX. More than 2/3^rd^ of the total 160 significant interactors belong to various known rRNA interacting families of proteins, or novel factors with a reported function in biogenesis. Proteins were color-coded as described in (B). **(D)** Interaction network analysis of *45S*-bound proteins identified by TREX, using the STRING physical interactions database (Szklarczyk *et al*., 2015). 153 out of the total 160 identified proteins can physically interact based on STRING analysis. Proteins were color-coded as described in (B).

### TREX defines the direct interactomes of *18S, 5.8S*, and *28S* rRNAs

Having validated the capability of TREX to comprehensively identify the proteins that directly interact with *45S* rRNA, we next set out to elucidate a detailed region-by-region interactome of *45S* rRNA, using the ability of TREX to define region-specific RNA-RBP interactions. First, we focused on mapping the interactomes of the *18S, 5.8S* and *28S* rRNA regions by selecting tiling oligonucleotides from the *45S* rRNA TREX that correspond to these segments (Fig. 4A). As before, the efficiency of each region-specific depletion was confirmed by RT-qPCR (Fig. S4A-C). Subsequently, we conducted quantitative MS analysis of the TREX samples originating from 10 million HCT116 cells per replicate experiment. Consistent with the pivotal role of the *18S* rRNA in constituting the 40S ribosomal subunit, we identified small ribosomal subunit proteins as the primary group of proteins that were directly bound to the *18S* rRNA (Fig. 4B, 4C, and Dataset S4). Notably, several known small subunit RAFs, including a number of translation initiation factors, SERBP1, LARP1, and EDF1, were among the significant interactors of *18S* rRNA (Fig. 4B). Moreover, we identified specific RBFs involved in small subunit biogenesis, such as RBM19, BYSL, TSR1, and LRRC47 (Dorner *et al*., 2023), among others (Fig. 4B). Interestingly, our analysis also revealed RPL24, a large ribosomal subunit protein, as a direct interactor of *18S* rRNA. This finding aligns with the resolved structure of the human 80S ribosome (Khatter et al., 2015), which demonstrated extensive contacts between the C-terminal portion of RPL24 and the *18S* rRNA post-translocation (Fig. S4D).

**Figure 4:**
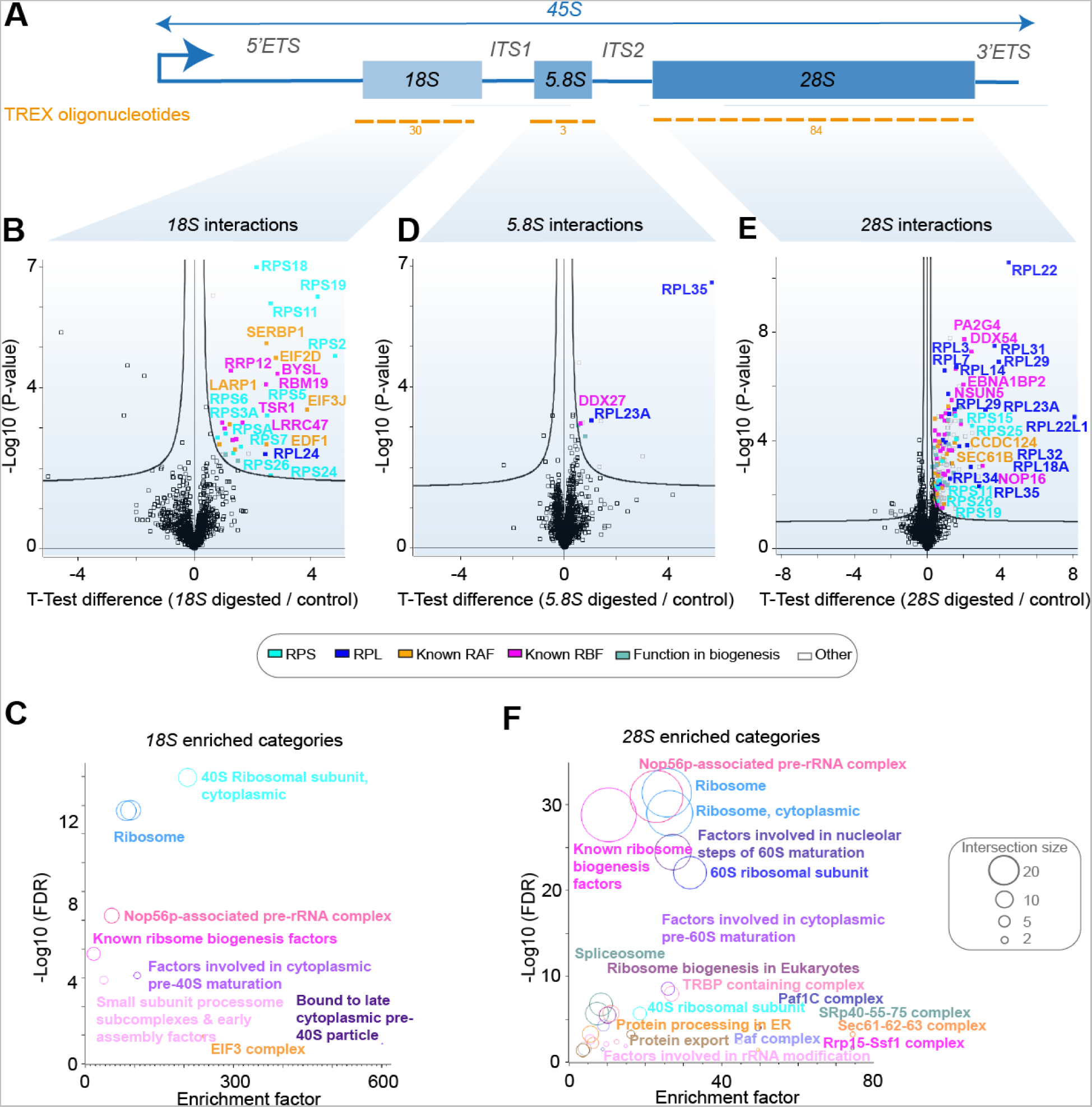
TREX defines the interactomes of *18S, 5.8S*, and *28S* rRNA. **(A)** Schematic representation of the *18S, 5.8S*, and *28S* segments within the *45S* rRNA. The tiling antisense DNA oligonucleotides used for depletion of each segment in each set of TREX experiments are depicted on the graph. **(B)** Volcano plot of the two-sample t-test comparison of *18S* rRNA digested vs undigested TREX samples, showing the enrichment of RPS proteins, known small subunit RAFs, known small subunit processing RBFs, as well as RPL24. Curved lines mark the significance boundary (FDR = 0.05, S0 = 0.1). Proteins were color-coded as described in (Fig. 3B). **(C)** Fisher’s exact test analysis of known protein categories that are over-represented amongst the *18S* interactors (Benjamini-Hochberg FDR < 0.05). Each circle represents an enriched category extracted from (Dorner *et al*., 2023) or Kyoto Encyclopedia of Genes and Genomes (KEGG) database (Kanehisa and Goto, 2000), with circle size representing the number of shared proteins. **(D)** Volcano plot of the two-sample t-test comparison of *5.8S* rRNA digested vs undigested TREX samples, showing the enrichment of DDX27, RPL23A, and RPL35. Curved lines mark the significance boundary (FDR = 0.05, S0 = 0.1). Proteins were color-coded as described in (Fig. 3B). **(E)** Volcano plot of the two-sample t-test comparison of *28S* rRNA digested vs undigested TREX samples, showing the enrichment of RPL proteins, known large subunit RAFs, known large subunit processing RBFs, as well as RPS11, RPS15, RPS19, RPS25, and RPS26. Curved lines mark the significance boundary (FDR = 0.05, S0 = 0.1). Proteins were color-coded as described in (Fig. 3B). **(F)** Fisher’s exact test analysis of known protein categories that are over-represented amongst the *28S* interactors (Benjamini-Hochberg FDR < 0.05). Each circle represents an enriched category extracted from (Dorner *et al*., 2023) or KEGG database, with circle size representing the number of shared proteins.

TREX analysis of *5.8S* rRNA demonstrated a notably smaller set of proteins directly interacting with this region (Fig. 4D and Dataset S5). Among them, we identified DDX27, a DEAD box helicase known for its specific involvement in 5.8S processing (Dorner *et al*., 2023). Additionally, RPL35 and RPL23A were found as the sole ribosomal proteins that significantly interacted with the *5.8S* rRNA (Fig. 4D). This is also in line with the resolved structure of the human ribosome (Khatter *et al*., 2015), in which RPL35 and RPL23A are the only ribosomal proteins that form extensive contacts with the *5.8S* rRNA (Fig. S4E), emphasizing the precision of TREX in revealing specific RNA-RBP interactions.

In contrast, TREX analysis of *28S* rRNA unveiled a multitude of direct interactors, primarily comprising large ribosomal subunit proteins (Fig. 4E, 4F, and Dataset S6). In addition, several RAFs that are known to interact with the large ribosomal subunit, including members of the translocon complex (Pfeffer et al., 2016) and CCDC124 (Wells et al., 2020), along with various large subunit RBFs such as PA2G4, NSUN5, DDX54, and EBNA1BP2 (Dorner *et al*., 2023), were among the identified proteins (Fig. 4E, 4F, and Dataset S6). Our analysis also revealed the presence of several small ribosomal subunit proteins. However, most of these proteins were observed to localize to the interface between the small and large ribosomal subunits in the human ribosome structure, where they can directly contact the *28S* rRNA (Fig. S4F). Interestingly, our TREX analysis also revealed evidence of existing ribosome heterogeneity in HCT116 cells, generated by incorporation of ribosomal protein isoforms. Specifically, RPL22 and RPL22L1 were both found to interact with *28S* rRNA (Fig. S4G), indicative of their successful incorporation into the large ribosomal subunits of HCT116 cells. In contrast, although both RPL7 and RPL7L were expressed in these cells, only RPL7 was significantly associated with *28S* rRNA, suggesting that RPL7L is unlikely to be contributing towards generation of ribosome heterogeneity in these cells (Fig. S4G). Taken together, these results characterize the direct interactomes of *18S, 5.8S*, and *28S* rRNAs in living cells, highlighting the robustness of TREX for comprehensive yet accurate profiling of distinct rRNA-protein interactions.

### TREX reveals the direct interactomes of *5’ETS, ITS1, ITS2,* and *3’ETS* spacer regions

Having revealed the direct binders of *18S, 5.8S*, and *28S* rRNA segments, we next interrogated the spacer regions of the *45S* pre-rRNA, by selecting tiling oligonucleotides from the *45S* rRNA TREX that correspond to these segments (Fig. 5A). As with other regions of *45S* rRNA, we validated the efficiency of each region-specific depletion through RT-qPCR (Fig. S5A-D). Subsequently, we conducted quantitative MS analysis of the TREX samples originating from 10 million HCT116 cells per replicate experiment. TREX analysis of the *5’ETS* revealed a predominant interaction with known RBFs involved in early pre-rRNA processing. For instance, several UTP complex members such as PWP2, DDX21, DCAF13, NOL11, UTP14a, UTP15, UTP18, WDR46, and WDR75, which are known to directly bind to *5’ETS* and mediate subsequent cleavage and processing events (Dorner *et al*., 2023), were amongst the prominent interactors (Fig. 5B, 5C, and Dataset S7). Moreover, a number of other key interactors of the *5’ETS* such as NCL (Ginisty et al., 1998), and DNTTIP2 (Rempola et al., 2006), were similarly identified (Fig. 5B and Dataset S7). As expected, ribosomal proteins were not prominently represented among the identified interactors of the *5’ETS*. However, three ribosomal proteins, namely RPS9, RPS25, and RPL23A, were still observed to interact with the *5’ETS* (Fig. 5B and Dataset S7). While RPS25 and RPL23A were also found to interact with other spacer regions, indicating potential roles in ribosome biogenesis beyond their canonical ribosomal functions, RPS9 specifically emerged as a *5’ETS* interactor. Inspection of the human 80S ribosome structure (Khatter *et al*., 2015) revealed a direct association of RPS9 with the 5’ end of *18S* rRNA (Fig. S5E), where the junction between the *5’ETS* and *18S* segment (cleavage site 1) once stood. Consequently, the release of RPS9 in *5’ETS* TREX may be attributed to its binding across this junction site prior to the cleavage.

**Figure 5:**
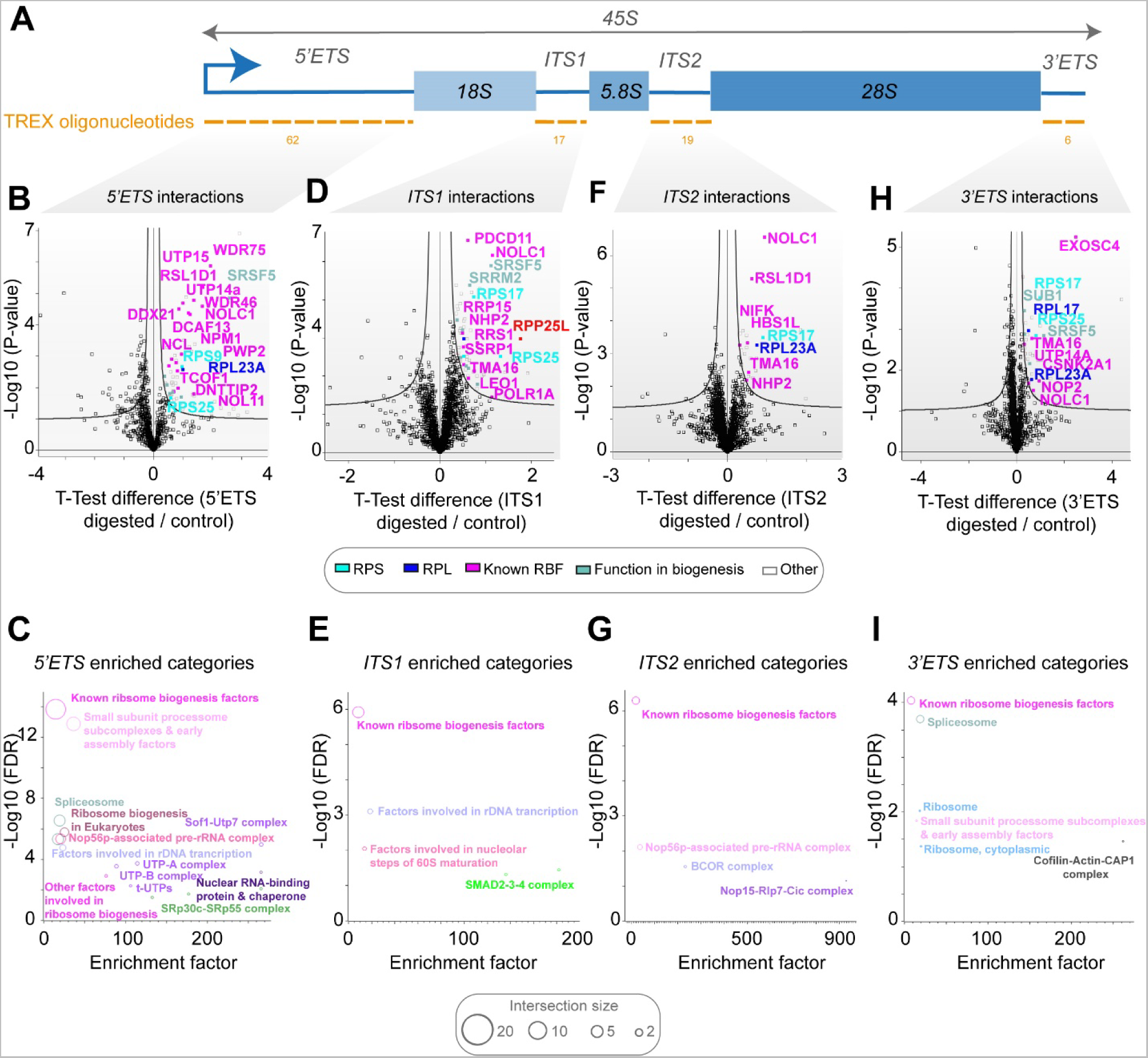
TREX defines the interactomes of *5’ETS*, *ITS1, ITS2*, and *3’ETS* spacer regions. **(A)** Schematic representation of the *5’ETS, ITS1, ITS2*, and *3’ETS* segments within the *45S* rRNA. The tiling antisense DNA oligonucleotides used for depletion of each segment in each set of TREX experiments are depicted on the graph. **(B)** Volcano plot of the two-sample t-test comparison of *5’ETS* digested vs undigested TREX samples, showing the prominent enrichment of several known early processing RBFs, along with RPS9, RPS25, and RPL23A, and SRSF5. Curved lines mark the significance boundary (FDR = 0.05, S0 = 0.1). Proteins were color-coded as described in (Fig. 3B). **(C)** Fisher’s exact test analysis of known protein categories that are over-represented amongst the *5’ETS* interacting proteins (Benjamini-Hochberg FDR < 0.05). Each circle represents an enriched category extracted from (Dorner *et al*., 2023) or KEGG database, with circle size representing the number of shared proteins. **(D)** Volcano plot of the two-sample t-test comparison of *ITS1* digested vs undigested TREX samples, showing the enrichment of various RBFs, RPS17, RPS25, SRSF5, SRRM2, and RPP25L. Curved lines mark the significance boundary (FDR = 0.05, S0 = 0.1). Proteins were color-coded as described in (Fig. 3B). **(E)** Fisher’s exact test analysis of known protein categories that are over-represented amongst the *5’ETS* interacting proteins (Benjamini-Hochberg FDR < 0.05). Each circle represents an enriched category extracted from (Dorner *et al*., 2023) or KEGG database, with circle size representing the number of shared proteins. **(F)** Volcano plot of the two-sample t-test comparison of *ITS2* digested vs undigested TREX samples, showing the enrichment of several known RBFs, along with RSP17 and RPL23A proteins. Curved lines mark the significance boundary (FDR = 0.05, S0 = 0.1). Proteins were color-coded as described in (Fig. 3B). **(G)** Fisher’s exact test analysis of known protein categories that are over-represented amongst the *ITS2* interacting proteins (Benjamini-Hochberg FDR < 0.05). Each circle represents an enriched category extracted from (Dorner *et al*., 2023) or KEGG database, with circle size representing the number of shared proteins. **(H)** Volcano plot of the two-sample t-test comparison of *3’ETS* digested vs undigested TREX samples, showing the enrichment of EXOSC4 and several other known RBFs, along with SUB1, SRSF5, RSP17, RPS25, RPL23A, and RPL17. Curved lines mark the significance boundary (FDR = 0.05, S0 = 0.1). Proteins were color-coded as described in (Fig. 3B). **(I)** Fisher’s exact test analysis of known protein categories that are over-represented amongst the *3’ETS* interacting proteins (Benjamini-Hochberg FDR < 0.05). Each circle represents an enriched category extracted from (Dorner *et al*., 2023) or KEGG database, with circle size representing the number of shared proteins.

TREX analysis of *ITS1 and ITS2* also revealed several known RBFs, but these were mostly distinct from the *5’ETS* interactors (Fig. 5D-G, Datasets S8 and S9). For instance, we identified RRP15 and PDCD11 amongst the identified interactors of *ITS1*, both of which are known to be important for *ITS1* processing (Dorner *et al*., 2023). Additionally, a prominently enriched protein among the identified *ITS1* interactors was RPP25L (Fig. 5D and Dataset S8), a paralogue of RPP25, which constitutes a subunit of RMRP endonuclease complex (Perederina et al., 2020). This ribonucleoprotein complex facilitates the critical site 2 cleavage within the *ITS1*, separating the pre-40S and pre-60S processing particles during ribosome biogenesis (Goldfarb and Cech, 2017). Remarkably, the protein expression level of RPP25L in HCT116 cells was found to be over 20-fold higher than that of RPP25, indicating a likely expression-based switch between the two paralogues in these cells (Fig. S5F). Amongst the direct interactors of *ITS2*, we found NIFK, the mammalian homologue of Nop15 that plays a key role in mediating *ITS2* folding (Fig. 5F and Dataset S9). Additionally, TMA16 and NHP2, two RBFs involved in pre-60S processing and modification (Ismail et al., 2022; Liang et al., 2020), were found to be direct binding partners of both ITS regions (Fig. 5D, 5F, and Datasets S8 & S9). Our investigation also revealed a previously unknown interaction between *ITS1* and SMAD family of proteins, which are traditionally associated with TGFB signaling (Fig. 5E, and Dataset S8). This discovery aligns with a recent study that unveiled an unanticipated role for *Drosophila* smad2 in regulating ribosome biogenesis at the *ITS1* cleavage stage (Martins et al., 2017), thereby supporting a potentially direct functional association between SMADs and *ITS1* in human cells as well.

Lastly, we examined the interactome of the *3’ETS*, the terminal transcribed section of the *45S* pre-rRNA, which has remained poorly characterized, compared to other spacer regions in mammalian cells. We identified EXOSC4, a core subunit of the exosome complex, as one of the top interactors of *3’ETS* (Fig. 5H and Dataset S10). This is consistent with recent evidence that shows *3’ ETS* containing pre-rRNAs can recruit the exosome complex for pre-rRNA surveillance, providing further support for a direct association (Shan et al., 2023). In addition to EXOSC4, several other RBFs were also identified as significant direct interactors of *3’ETS* (Fig. 5H, 5I and Dataset S10). These included pre-60S processing factors such as TMA16 and NOP2, as well as early processing factors such as UTP14a (Fig. 5H, 5I and Dataset S10), the latter supporting a potential coupling of certain *5’ETS* and *3’ETS* processing events, which has been suggested to occur in human cells (Mullineux and Lafontaine, 2012). Similar to the other spacer regions, ribosomal proteins were not prominently identified amongst the *3’ETS* interactors, but a select number of ribosomal proteins were still present. Of these, RPS17, RPS25, and RPL23A were also identified as interactors of other spacer regions, whereas RPL17 exhibited specificity for the *3’ETS* (Fig. 5H and Dataset S10). Similar to the case of RPS9 and the *5’ETS*, we could demonstrate that the direct contact site for RPL17 in the human 80S ribosome structure is formed at the 3’ end of *28S* rRNA, where the junction with the *3’ETS* (cleavage site 02) was once located (Fig. S5G). Consequently, the release of RPL17 in our TREX analysis of *3’ETS* may be attributed to its binding across this junction site prior to its cleavage. Collectively, these TREX findings provide, for the first time, a comprehensive snapshot of the direct interactomes associated with each transcribed spacer region of the human *45S* pre-rRNA under endogenous conditions.

### Cross-comparison of TREX interactomes reveals region-specific and multi-regional interactors that regulate ribosome biogenesis

Next, we set out to integrate the TREX results obtained from different *45S* rRNA segments, in order to map the positional landscape of the identified RNA-protein interactions across the full length of *45S* pre-rRNA. For this purpose, we selected proteins exhibiting significant interactions with at least one segment of *45S* rRNA, and conducted unsupervised hierarchical clustering of the t-test scores across the different TREX experiments. Remarkably, more than 95% of the interacting proteins (381 out of 400) displayed highly specific positional interaction patterns, forming distinct clusters primarily associated with a specific segment of *45S* pre-rRNA (Fig. 6A and Dataset S11). In particular, clusters 1 and 2 comprised interactors specific to the *18S* segment, primarily encompassing ribosomal small subunit proteins (RSPs) and small subunit RBFs. Cluster 3 consisted of interactors specific to the *5’ETS*, prominently featuring members of different UTP complexes. Interactors specific to *ITS1* were in cluster 4, featuring known participants such as PDCD11 and RPP25L. Cluster 5 encompassed interactors of both the *5.8S* and *ITS2* regions, featuring established partners such as NIFK. Clusters 6 and 7 primarily constituted interactors specific to the *3’ETS*, while clusters 8 and 10, comprising the largest group of region-specific interactors, were predominantly associated with the *28S* segment, containing many large subunit RBFs and ribosomal large subunit proteins (RPLs) (Fig. 6A and Dataset S11). RPL35, was the only protein that did not cluster with any other interactor, as it exhibited a unique profile of major association with *5.8S*, with some binding also to *28S* (Dataset S11), a pattern consistent with its above mentioned positioning in the human 80S ribosome structure (Fig. S4E).

**Figure 6:**
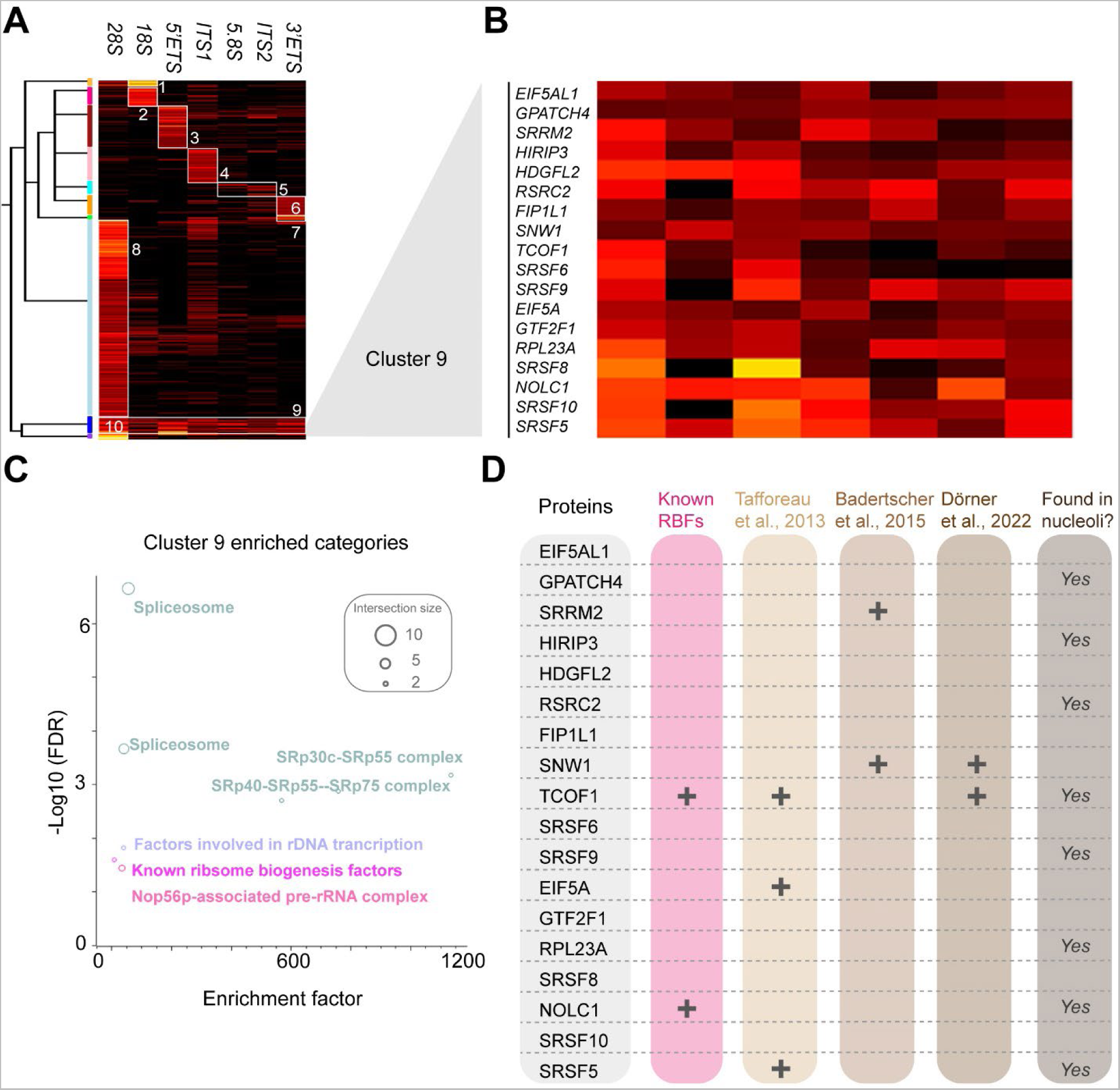
TREX reveals region-specific and multi-regional interactors of *45S* rRNA. **(A)** Unsupervised Hierarchical clustering analysis of t-scores for significant hits from TREX analyses of different segments of *45S* rRNA, with complete Euclidean distance calculation and K-means pre-processing (t-score color scale: black = 0 or less; red = 2-6; orange = 6-10; yellow = 10 or more). The majority of identified proteins separate into nine clusters (clusters 1, 2, 3, 4, 5, 6, 7, 8, and 10) that exhibit region-specific patterns of binding to individual *45S* rRNA segments, while members of cluster 9 exhibit binding to several segments. **(B)** Zoomed-in view of cluster 9 and the list of its constituent proteins. **(C)** Fisher’s exact test analysis of known protein categories that are over-represented amongst the cluster 9 proteins (Benjamini-Hochberg FDR < 0.05). Each circle represents an enriched category extracted from (Dorner *et al*., 2023) or KEGG database, with circle size representing the number of shared proteins. **(D)** Overlap analysis of cluster 9 proteins with lists of known RBFs from (Dorner *et al*., 2023) and hits from three large-scale RNAi screens of human ribosome biogenesis regulators (Badertscher *et al*., 2015; Dorner *et al*., 2022; Tafforeau *et al*., 2013), reveals 6 out of the total 18 proteins to have a role in ribosome biogenesis. Proteins with reported localization to the nucleolus, either as primary or as an additional site of localization, according to the Human Protein Database, are also marked on the graph.

In contrast to the majority of identified clusters that were highly region-specific, cluster 9 exhibited enrichment across multiple regions (Fig. 6A and Dataset S11), suggesting a distinct mode of interaction. Cluster 9 comprised 18 proteins, including NOLC1 and TCOF1, two RBFs known to form a complex that couples rDNA transcription with the rRNA modification machinery (Werner et al., 2015), as well as several splicing factors such as SNW1, SRSF5, SRSF6, SRSF9, SRSF10, SRRM2, and RSRC2 (Fig. 6B, 6C, and Dataset S11). Previous studies have implicated certain Serine/Arginine (SR)-rich splicing factors such as SRSF1 and PRP43 in regulation of various pre-rRNA-related processes (Aquino et al., 2021; Bohnsack et al., 2009; Kress et al., 2008). However, our findings unveil a more extensive repertoire of splicing factors directly associating with multiple pre-rRNA regions, suggesting a prominent moonlighting function for this family of proteins in ribosome biogenesis. In agreement with this, five members of cluster 9 have been identified as hits in the above-mentioned large-scale functional RNAi screens for human ribosome biogenesis regulators (Badertscher *et al*., 2015; Dorner *et al*., 2022; Tafforeau *et al*., 2013), supporting their functional involvement in pre-rRNA related processes (Fig. 6D). Moreover, subcellular localization analysis using the Human Protein Atlas database (Thul et al., 2017), revealed that eight members of the cluster 9 can localize to the nucleolus, further supporting an involvement in rRNA related processes and regulation of ribosome biogenesis (Fig. 6D). Collectively, our analysis depicts a specific positional binding pattern for the majority of *45S* rRNA-associated proteins, but we also identify a distinct cluster of proteins that include several splicing factors, which interact with multiple pre-rRNA regions. Such direct association with multiple pre-rRNA segments, combined with the functional impact of their depletion, and a nucleolar subcellular localization, is indicative of a novel role for these factors in regulation of ribosome biogenesis.

## Discussion

The identification and characterization of proteins that interact with specific RNA sequences within living cells is crucial for understanding the intricate mechanisms that govern RNA-mediated processes. In this study, we present TREX, a highly efficient and specific RNA-centric method that fills a critical gap in the field by enabling unbiased and region-specific exploration of direct RNA-protein interactions in their endogenous context. The power of TREX lies in its innovative combination of phase extraction and RNase H-mediated RNA degradation (Fig. 1A). Leveraging the highly efficient and specific capabilities of phase extraction, TREX effectively separates RNA-bound from non-bound proteins. This, combined with RNase H-mediated degradation, which robustly and selectively cleaves RNA in RNA-DNA hybrids, enables the precise release and recovery of proteins that were bound to a given RNA target sequence. The unique integration of phase extraction and RNase H-mediated degradation establishes TREX as a powerful method for region-specific investigation of RNA-protein interactions under endogenous settings in living cells. In fact, the only currently available RNA-centric approach for assessing region-specific RNA-RBP interactions in cells is incPRINT, which uses a reporter-based over-expression system in which tagged RBPs and RNA sequences of interest are ectopically co-expressed in a cell-line, followed by cell lysis and assessment of RNA-RBP co-purification via a luminescence based reporter assay (Graindorge et al., 2019). In comparison, TREX is unbiased rather than candidate based, works with any region of any endogenous RNA, does not require any genetic manipulations and RNA/protein tagging, and allows the interactome profiling of any target sequence of interest with minimal preparations. An additional advantage of TREX lies in its remarkable cost-effectiveness, as it solely requires unlabeled tiling antisense DNA oligonucleotides, along with generic reagents commonly used in most RNA biology laboratories such as TRIZOL. This affordability significantly enhances the accessibility of TREX to researchers worldwide, empowering them to explore the direct binding partners of any regions of any target RNA, in diverse biological contexts.

RNA affinity capture is currently the most commonly used approach for analysis of direct protein interactors of a given RNA target under endogenous settings (Hafner *et al*., 2021). However, methods based on this approach are unsuited for region-specific interaction analysis, as a target RNA molecule can only be purified in its entirety through affinity capture. Moreover, RNA affinity capture based methods often suffer from low efficiency of protein-recovery, necessitating large quantities of input material in order to obtain meaningful MS results (McHugh *et al*., 2015; McHugh and Guttman, 2018). TREX overcomes both these shortcomings through the combined use of organic phase extraction and RNase H mediated degradation. The robust enzymatic reaction of RNase H enables near complete region-specific target degradation, and stoichiometric release of any associated proteins, with phase extraction providing an immaculate system to separate the RNA-bound from the released RBPs, thus greatly boosting both the specificity and efficiency of extractions. To assess the performance of TREX in comparison with RNA affinity capture based methods, we conducted benchmarking experiments using the well-characterized *U1* snRNA as the target. Our results demonstrated that with as low as 5 to 10% of the starting input material, TREX could achieve superior performance compared to three previous affinity capture-based studies, in terms of revealing known *U1* binding partners. We also demonstrated the ability of TREX to investigate region-specific interactions, by applying it to reveal the direct binding partners of the most conserved segment of *NORAD* lncRNA, a non-coding RNA with key functions in maintenance of genome stability (Lee *et al*., 2016; Tichon *et al*., 2016). Our analysis showed a substantial overlap between TREX and previous *NORAD* interactome capture studies, revealing several known binding partners such as PUM1, SAM68, IGF2BP2/3, PABPN1 and RBMX. Crucially, we also found many previously unknown interactors. Amongst these, TOP1 was found as one of the most significant hits, a finding that is strongly in line with the reported functional interplay between in the maintenance of genome stability (Munschauer *et al*., 2018).

Leveraging the power of TREX for deciphering region-specific RNA-protein interactions in an RNA-centric manner, we generated a comprehensive region-by-region interactome map of human *45S* rRNA. This revealed a highly specific interactome landscape for each segment of *45S* rRNA, which was consisted of specific sets of ribosomal proteins, RBFs, RAFs, as well as previously unknown rRNA-binding proteins, some of which were shown to be functionally important for ribosome biogenesis. Remarkably, while most of the identified proteins exhibited region-specific binding profiles, our analysis unveiled a surprising cohort of proteins that were bound to multiple *45S* rRNA regions. Several splicing factors were present amongst this group, some of which are known to impact ribosome biogenesis upon depletion. Multiple members also localized to the nucleolus, a feature strongly linked to ribosome biogenesis. Together, these results reveal a previously unknown group of multi-regional pre-rRNA interactors that are involved in regulation of ribosome biogenesis. The exact function and mechanism of action of these proteins in the context of ribosome synthesis remains to be determined.

Transcribed in the nucleolus by RNA polymerase I (RNAPI), *45S* rRNA is a large polycistronic transcript consisted of *18S*, *5.8S*, and *28S* rRNA sequences that form the RNA core of the large and small ribosomal subunits. These sequences are interspersed with *5’ETS*, *ITS1, ITS2* and *3’ ETS* spacer sequences, which are ultimately cleaved and removed from maturing rRNA during the process of ribosome biogenesis. It is now clear that as nascent *45S* rRNA emerges from RNAPI, it becomes rapidly decorated by several ribosomal proteins and RBFs, giving rise to a distinct nucleolar particle known as the 90S pre-ribosome. Through a series of endonucleolytic cleavages and exonucleolytic processing events, along with rRNA modifications, folding, and assembly of additional ribosomal proteins, RBFs, and *5S* rRNA, the 90S particle splits into pre-40S and pre-60S particles, and gradually matures into the translationally competent small and large ribosomal subunits (Dorner *et al*., 2023). Critically, recent advances in determining the composition and structures of ribosome biogenesis intermediates have primarily relied on affinity purification of selected RBFs that associate with specific stages of ribosome biogenesis (Bassler and Hurt, 2019; Fatica and Tollervey, 2002). Consequently, the emerging understanding of ribosome biogenesis intermediates that is derived from such studies is primarily protein-centric, with interactions/intermediates that may not be particularly amenable to survive protein affinity purifications likely to be under-represented or lost. Our TREX analysis, on the other hand, provides an alternative RNA-centric view of the composition of factors associated with each segment of *45S* rRNA, thus revealing potentially novel ribosome biogenesis components that could have been missed by protein-centric approaches.

In conclusion, TREX is a transformative method for RNA-centric profiling of RNA-protein interactions in living cells. The combination of efficiency, specificity, and region-specific mapping capability positions TREX as a promising tool for investigating RNA-protein interactions in living cells. Moreover, due to its ease of implementation and its applicability to any cell-type or RNA of interest, TREX can be easily adopted and used for diverse types of studies that are aimed at assessing RNA-protein interactions from an RNA-centric perspective.

## Experimental procedures

Full details of the TREX tiling antisense DNA oligonucleotides used in this study (Table S1), as well as the RT-qPCR oligonucleotide probes (Table S2), are available in the **Supplementary Information**.

For a detailed step-by-step protocol of TREX, please visit: https://www.mardakhehlab.info/resources/trex

### Key resource table

| REAGENT | SUPPLIER | CATALOGUE NUMBER |
| --- | --- | --- |
| **TREX** |  |  |
| Invitrogen TURBO DNase and 10X Reaction Buffer | Fisher Scientific UK Ltd | 10646175 |
| UltraPure 10% SDS | Life Technologies Ltd | 15553-035 |
| Thermostable RNase H and RNase H Reaction Buffer | New England Biolabs UK Ltd | M0523S |
| Ambion RNase H | Fisher Scientific UK Ltd | AM2293 |
| RNasin Plus RNase Inhibitor | Promega Ltd | N2611 |
| Invitrogen Nuclease-Free Water | Fisher Scientific UK Ltd | 10526945 |
| 1M Tris pH 7.0 | Thermo Fisher Scientific | AM9850G |
| 0.5M EDTA pH 8.0 | Thermo Fisher Scientific | AM9260G |
| 1M MgCl<sub>2</sub> | Thermo Fisher Scientific | AM9530G |
| Isopropanol | Fisher Scientific UK Ltd | P/7507/PB17 |
| Ethanol | Fisher Scientific UK Ltd | E/0650DF/17 |
| Acetone | Merck | 34850-2.5L |
| Chloroform | Honeywell | 650471-1L |
| 5M NaCl | Thermo Fisher Scientific | AM9760G |
| TRI REAGENT LS (TRIZOL LS) | Sigma-Aldrich | T3934-200ML |
| TRI REAGENT (TRIZOL) | Sigma-Aldrich | 93289-25ML |
| Nuclease-Free Water | Fisher | 10526945 |
| **MS sample preparation** |  |  |
| Urea | Sigma-Aldrich | U1250-5KG |
| Ammonium bicarbonate | Sigma-Aldrich | A6141-500G |
| Thermo Scientific Pierce MS Grade Trypsin Protease | Fisher Scientific UK Ltd | 13474189 |
| Iodoacetamide | Biosciences | 786-228 |
| DTT, molecular biology grade (Dithiothreitol, Cleland's reagent) | Thermo Scientific | R0861 |
| Vivacon 500, 30,000 MWCO Hydrosart | Sartorius | VN01H22 |
| Pierce High pH Reversed-Phase Peptide Fractionation Kit | Life Technologies | 84868 |
| Empore™ Octadecyl C18 47mm Extraction Disks 2215 | Empore™ | 66883-U |
| TFA, LC-MS Grade | Thermo Scientific | 85183 |
| Acetonitrile HPLC Far UV/ Gradient Grade | J.T.Baker | 9012 |
| Acetic Acid Glacial | Scientific Laboratory Supplies | 200-580-7 |
| Trypsin | Merck | T6567-1MG |
| **Library preparation and sequencing** |  |  |
| PCR Add-on Kit for Illumina | Lexogen | 020 |
| RiboCop rRNA Depletion HMR V2 kit for ribosomal RNA + CORALL Total RNA-seq V2 kit | Lexogen | 183-184 |
| UDI 12 nt set B1 | Lexogen | UDI12B_0001-0096 |
| **RT-qPCR** |  |  |
| EDTA | Thermo | 15575-038 |
| Proteinase K | Thermo | 25530049 |
| Power SYBR Green RNA-To-Ct 1 Step kit | Applied Biosystems | 4389986 |
| SDS | Thermo | 15553027 |
| TRI Reagent LS | Sigma-Aldrich | T3934 |
| Tris HCl pH 8.0 | Thermo | AM9855G |
| **Tissue culture** |  |  |
| HCT116 colon carcinoma cells | ATCC | CCL-247 |
| McCoy's 5A (Modified) Medium, GlutaMAX™ Supplement | Thermo | 36600088 |
| Trypsin/EDTA solution | Fisher Scientific | R-001-100 |
| Gibco Fetal Bovine Serum | Thermo Fisher | 10437028 |
| **Commercial kits and reagent sets** |  |  |
| Pierce™ BCA Protein Assay Kit | Thermo Fisher | 23225 |
| MycoAlert™ PLUS Mycoplasma Detection Kit | Lonza | LT07-705 |

### Cell culture

HCT116 cells were cultured in McCoy’s 5a medium supplemented with 10% fetal bovine serum (FBS) and 100 U of penicillin/0.1 mg mL−1 streptomycin. The cells were maintained at 37°C in a humidified incubator with 5% CO_2_. HCT116 were routinely examined for mycoplasma contamination and their identity was confirmed by short tandem repeat profiling.

### TREX

HCT116 cells were seeded on 150 mm TC-treated culture dishes and allowed to attach for a minimum of 16 hours, prior to being washed with ice-cold PBS and irradiated on ice with 200 mJ/cm^2^ of UV-C (254 nm), using a Hoefer Scientific UV Crosslinker. Cells were then lysed by direct addition of TRIZOL to each dish (1 ml per ~20 million cells). We used 100 million cells per biological replicate experiment for the *NORAD* ND4 TREX analysis, while 10 million cells per replicate were used for all other TREX experiments. After scraping the cells in TRIZOL, homogenized lysates were incubated for 5 min at room temperature to dissociate non-crosslinked RNA-protein interactions, before the addition of chloroform (200 µl per 1 ml of lysate) and centrifugation for 15 min at 12,000 g at 4°C to induce phase separation. The aqueous and organic phases were subsequently discarded, and the interface was resolubilized in TRIZOL. Phase separation and resolubilization was repeated two more times to remove any non-crosslinked RNA or proteins. The isolated interface was then gently washed with TE buffer to remove any traces of TRIZOL, before being solubilized by resuspension in TE buffer supplemented with increasing concentrations of SDS, as described previously (Trendel *et al*., 2019). The RNA-protein crosslinks were then precipitated by adding NaCl (to 300 mM final concentration) followed by isopropanol (to 50% final concentration), and the pellet was washed with 70% ethanol, and resuspended in nuclease-free water. Any contaminating genomic DNA was removed by treating the samples with TURBO DNase (2U per million cells) for 50 min at 37 °C, followed by a further round of isopropanol precipitation. The pellets were washed again with 70% ethanol and resuspended in probe hybridization buffer (50mM NaCl, 1mM EDTA, 100mM Tris HCl pH 7.0), before a pool of tiling antisense DNA oligonucleotides complementary to the target RNA region of interest was added to the mixture. The individual oligonucleotides used were non-overlapping, on average 60 nt long (and no shorter than 30 or longer than 90), and unmodified. We ensured that no oligonucleotides strongly matched to an off-target RNA by blasting their sequences against the RefSeq database. The sequences of all TREX antisense tiling probes used in this study can be found in Table S1. The amount of oligo pools used per reaction was determined prior to the TREX experiment by performing oligo dose titration for each target, followed by RT-qPCR analysis (as shown in Fig. S1). Annealing was performed in a Thermomixer by first heating the samples to 95 °C for 2 min to denature all RNAs, followed by a gradient temperature drop of 2°C per minute, until the samples reached 50°C. DNA-hybridized RNA regions were subsequently digested following the addition of MgCl_2_ to neutralize the EDTA, followed by Thermostable RNase H (1U per million cells) to each reaction and further incubation at 50°C for 60 min. As control, samples were treated identically but no RNase H enzyme was added. Samples were then centrifuged at 12000 g for 3 min and the supernatant was transferred to a new tube. 10% of each supernatant was aliquoted and taken for RNA extraction to analyze target depletion efficiency and specificity (see below), whilst the remaining 90% was subjected to a second round of organic phase separation by adding 900µl of TRIZOL LS to each sample, followed by 200 µl of chloroform, mixing, and centrifugation for 15 min at 12,000 g at 4°C. The released RBPs were recovered by taking the organic phase into a new tube. Ice-cold acetone was added to each sample (final concentration of 80%), followed by mixing and overnight incubation at −20°C in order to precipitate the released RBPs. The pellets were subsequently recovered and washed with 1 ml of 80% acetone, followed by centrifugation for 15 min at 16,000 g at 4°C. This step was repeated once, and the final pellet was air-dried for 5 min at room temperature.

### Total lysate preparation

For label-free absolute protein quantification analysis of the HCT116 proteome, 500 µl aliquots of TRIZOL lysed HCT116 cells (corresponding to 10 million cells) prior to TREX analysis were taken and subjected to protein precipitation by adding ice-cold acetone to the final concentration of 80%, followed by mixing and overnight incubation at −20°C. The precipitated protein pellets were subsequently recovered and washed with 1 ml of 80% acetone, followed by centrifugation for 15 min at 16,000 g at 4°C. This step was repeated once, and the final pellet was air-dried for 5 min at room temperature.

### Mass spectrometry sample preparation and data acquisition

Acetone precipitated proteins from TREX were subjected to in-solution digestion by Trypsin as described previously (Azman et al., 2023). Briefly, proteins were recovered in 200 μl of 2M Urea, 50mM Ammonium Bicarbonate (ABC) and reduced by adding DTT to a final concentration of 10 mM. After 30 minutes of incubation at RT, samples were alkylated by adding 55 mM iodoacetamide and Incubation for another 30 minutes at RT in the dark. Trypsin digestion was then performed using 2 µg of trypsin per sample. The next day, samples were desalted using the Stage Tip procedure (Rappsilber et al., 2003), and recovered in 0.1% TFA, 0.5% Acetic Acid, 2% Acetonitrile (A* buffer) for LC-MS/MS analysis. For total proteomics analysis, samples were similarly dissolved in 200 μl of 2M Urea, 50mM Ammonium Bicarbonate (ABC), reduced and alkylated with DTT and iodoacetamide, followed by trypsin digestion of an equivalent of ~100 μg of protein, using the FASP protocol (Wisniewski et al., 2009). The digested peptides were subjected to peptide fractionation using Pierce™ High pH reverse-phase fractionation kit, as described previously (Dermit et al., 2020). Fractions were dried with vacuum centrifugation before recovery in A* buffer for LC-MS/MS analysis. LC-MS/MS analysis was performed as described before (Azman *et al*., 2023), using a Q Exactive-plus Orbitrap mass spectrometer coupled with a nanoflow ultimate 3000 RSL nano HPLC platform (Thermo Fisher). For total proteomics analysis, equivalent of ~1g of total protein per fraction was injected into the instrument, while ~90% of the total peptide mixture per each TREX experiment was injected. Samples were resolved at flow rate of 250 nL/min on an Easy-Spray 50cm X 75 m RSLC C18 column (Thermo Fisher), using a 123 min gradient of 3% to 35 % of Buffer B (0.1% FA in Acetonitrile) against Buffer A (0.1% FA in LC-MS gradient water). LC-separated samples were infused into the MS by electrospray ionization (ESI). Spray voltage was set at 1.95 kV, and capillary temperature was set to 255°C. MS was operated in data dependent positive mode, with 1 MS scan followed by 15 MS2 scans (top 15 method). Full scan survey spectra (m/z 375-1,500) were acquired with a 70,000 resolution for MS scans and 17,500 for the MS2 scans. A 30 sec dynamic exclusion was applied to all runs.

### Mass spectrometry data analysis

Maxquant (version 1.6.3.3) was used for all mass spectrometry search and quantifications (Tyanova et al., 2016a). Raw data files were searched against a FASTA file of the human proteome, excluding variants and isoforms, extracted from UNIPROT (2023). Enzyme specificity was set to “Trypsin”, allowing up to two missed cleavages. Label-free quantification was enabled using LFQ intensity calculation with a minimum ratio count of two. Variable modifications were set at Oxidation (M), Acetylation (N-term), and Phosphorylation (ST). False discovery rates (FDR) were calculated using a reverse database search approach, and was set at 1% for identification of peptides, modifications, and proteins. “Match between runs”, “Re-quantify”, and “iBAQ” options were enabled. Default Maxquant parameters were used for all other settings. All downstream data analyses such as data filtering, Log transformation, imputation of missing values (downshift = 1.8 s.d., variation = 0.3 s.d.), two-sample t-test analysis, category enrichment analysis, and hierarchical clustering, were performed by Perseus (version 1.6.2.3) (Tyanova et al., 2016b). PCA was performed on transformed and imputated data. Where one TREX sample behaved overtly differently to the other three or four biological replicates based on the PCA scattering, the outlier was subsequently removed from downstream analysis. For two-sample t-test analyses, permutation-based p-value correction with an FDR cut-off of < 0.05 and S0 of 0.1 was used. Category enrichment analysis was performed using Fisher’s exact test, with a Benjamini-Hochberg FDR cut-off of < 0.02. Clustering of t-scores, combined from different TREX t-test analyses, was performed using Euclidean complete distance calculation, with K-means preprocessing. Only proteins that were identified as significant binders in at least one TREX experiment were included for clustering analysis.

### RNA isolation, cDNA synthesis and qPCR

Target depletion efficiency upon RNAse H treatment was determined as following: an aliquot of solubilized and annealed interface, with or without RNase H treatment, corresponding to ~2 million cells was digested with 200 μg of Proteinase K in 200 μg of 10 mM Tris pH 8.0, 1 mM EDTA, 0.5% SDS for 1 hour at 56 °C. RNA was then extracted with TRIZOL LS, following the manufacturers’ instructions for RNA extraction, before cDNA synthesis and quantitative real-time PCR analysis in a single reaction, using the Power SYBR Green RNA-to-CT 1-Step kit. The QuantStudio 7 Flex real-time PCR system (Applied Biosystems) with the following program was employed for qPCR analysis: 48°C for 30 min, 95°C for 10 min followed by 40 cycles of 95°C for 15 s and 60 °C for 60 s. *GAPDH* and *RPS18* were used as housekeeping genes for input normalization, except for the *45S* region-specific experiments, where *18S* was used instead of *RPS18*, as to also provide a control for the region-specific depletions. RNA expression was estimated using 2^-ΔCT^. For *ITS2*, the qPCR probes used led to amplification of an additional non-specific peak with a lower melting temperature, so to delineate the specific peak from the non-specific one, relative area under the curve for the higher melting peak was specifically measured using the DescTools package in R. The sequences of all qPCR primers used in this study are provided in Table S2.

### RNA library preparation, sequencing and analysis

Libraries for RNA sequencing were prepared from 700 ng of purified RNA from RNase H treated or untreated TREX samples, using Lexogen RiboCop rRNA Depletion HMR V2 kit for ribosomal RNA depletion coupled to CORALL Total RNA-seq V2 kit. Lexogen UDI 12 nt was used as indexing system, and the PCR Add-on Kit for Illumina was used to optimize the number of PCR cycles. The PCR-amplified indexed libraries were sequenced to a depth of 40 million reads per sample, using 150bp paired end sequencing on an Illumina NovaSeq 6000 instrument (Novogene UK Company Limited, Cambridge). The quality of generated FASTQ files was checked with fastqc tool (version 0.11.9). Unique Molecular Identifiers (UMIs) were extracted from the initial FASTQ files using UMItools (version 1.1.1) (Smith et al., 2017). This involves the identification and removal of UMIs from each read, allowing for accurate quantification of gene expression and mitigation of amplification biases caused by PCR duplicates. The reads were subjected to trimming with trim-galore (version 0.6.5) to remove adapter sequences and low-quality reads with the following parameters –paired –retain_unpaired –illumina –gzip. Reads were aligned to the reference human genome GRCh38.p13 using the STAR aligner (version 2.7.9a) (Dobin et al., 2013). The alignment parameters were set as --outFilterType BySJout --outFilterMultimapNmax 20 --alignSJoverhangMin 8 --alignSJDBoverhangMin 1 -peOverlapNbasesMin 40 --peOverlapMMp 0.8 --outFilterMismatchNoverLmax 0.6 --alignIntronMin 20 --alignIntronMax 1000000 --alignMatesGapMax 1000000 --outSAMattributes NH HI NM MD --outSAMtype BAM SortedByCoordinate --quantMode TranscriptomeSAM --twopassMode Basic --outFilterScoreMinOverLread 0 --outFilterMatchNminOverLread 0 --outFilterMismatchNmax 2 -- limitOutSJcollapsed 2000000. The resulting alignment files were used to generate a matrix with counts and TPM values using the rsem-calculate-expression function from the RSEM software (version 1.3.1) (Li and Dewey, 2011), using the following options: --bam --strandedness forward --no-bam-output --paired-end. The TPM values were then Log2 transformed and filtered based on having valid values in at least 3 biological replicates in at least one sample group (RNase H treated or control), before imputation of missing values using the Perseus Imputation function (downshift = 1.8 s.d., variation = 0.3 s.d.). The values were then subjected to two-sample t-test analysis with permutation based FDR calculation (FDR <0.05 and S0 = 0.1) to reveal significantly different transcripts between the RNase H treated and untreated sets.

## Author contributions

<u>F.K.M.</u> conceived the study. <u>M.Dodel</u> optimized the methodology. <u>M.Dodel</u>, <u>G.G.</u>, and <u>S.K.</u> performed the experiments. <u>G.G.</u>, <u>M.Dermit,</u> and <u>F.K.M</u> analyzed the data. <u>F.K.M.</u> and <u>L.S.</u> acquired funding, supervised the work, and wrote the manuscript.

## Acknowledgements

We would like to acknowledge the BCI mass spectrometry core facility for their support with the proteomics experiments. We would also like to thank Kamil Kranc, Kevin Rouault-Pierre, Susana Godinho, Sarah McClelland, and Tyson Sharp for critical comments on the manuscript. This work was supported by Medical Research Council grants (MR/P009417/1 and MR/W001500/1) and Barts Charity grants (MGU0346 and G-002420) to <u>F.K.M</u>. <u>L.S.</u> is supported by Barts Charity (MGU0404), Cancer Research UK Career Establishment Award (RCCFEL\100007), Royal Society Research Grant (RGS\R1\231139), and the Academy of Medical Science Springboard Award (SBF006\1026). <u>G.G</u>. is supported by an AIRC Fellowship for Abroad.

## Declaration of interest

<u>F.K.M.</u>, <u>L.S.</u>, <u>M.Dermit</u>, <u>G.G.</u>, and <u>M.Dodel</u> are inventors and contributors on a pending patent covering the TREX method.

## Notes

https://www.mardakhehlab.info/resources/trex

### Competing Interest Statement

Faraz Mardakheh, Lovorka Stojic, Maria Dermit, Giulia Guiducci, and Martin Dodel are inventors and contributors on a pending patent covering the TREX method.

