## Supplementary Information for "TREX reveals proteins that bind to specific RNA regions in living cells"

Supplementary information for this manuscript consists of a supplementary excel file containing supplementary tables (Table S1-S2), a supplementary excel file containing supplementary datasets (Dataset S1-S11), and supplementary figures (Fig. S1-S5).

### Supplementary Tables

**Table S1: List of TREX antisense DNA oligonucleotides used in this study for RNase H-mediated depletion of each indicated target RNAs.**

**Table S2: List of all RT-qPCR primers used in this study.**

### Supplementary Datasets

**Dataset S1: List of identified proteins in the *U1* TREX experiment, with significantly interacting proteins marked by +. Significance was calculated using a permutation based FDR of < 0.05 and S0 of 0.1.**

**Dataset S2: List of identified proteins in the *NORAD* TREX experiment, with significantly interacting proteins marked by +. Significance was calculated using a permutation based FDR of < 0.05 and S0 of 0.1.**

**Dataset S3: List of identified proteins in the *45S* TREX experiment, with significantly interacting proteins marked by +. Significance was calculated using a permutation based FDR of < 0.05 and S0 of 0.1.**

**Dataset S4: List of identified proteins in the *18S* TREX experiment, with significantly interacting proteins marked by +. Significance was calculated using a permutation based FDR of < 0.05 and S0 of 0.1.**

**Dataset S5: List of identified proteins in the *5.8S* TREX experiment, with significantly interacting proteins marked by +. Significance was calculated using a permutation based FDR of < 0.05 and S0 of 0.1.**

**Dataset S6: List of identified proteins in the *28S* TREX experiment, with significantly interacting proteins marked by +. Significance was calculated using a permutation based FDR of < 0.05 and S0 of 0.1.**

**Dataset S7: List of identified proteins in the *5'ETS* TREX experiment, with significantly interacting proteins marked by +. Significance was calculated using a permutation based FDR of < 0.05 and S0 of 0.1.**

**Dataset S8: List of identified proteins in the *ITS1* TREX experiment, with significantly interacting proteins marked by +. Significance was calculated using a permutation based FDR of < 0.05 and S0 of 0.1.**

**Dataset S9: List of identified proteins in the *ITS2* TREX experiment, with significantly interacting proteins marked by +. Significance was calculated using a permutation based FDR of < 0.05 and S0 of 0.1.**

**Dataset S10: List of identified proteins in the *3'ETS* TREX experiment, with significantly interacting proteins marked by +. Significance was calculated using a permutation based FDR of < 0.05 and S0 of 0.1.**

**Dataset S11: T-test statistics scores for the combined list of significant interactors found each *45S* segment (datasets S4 to S10), and the clusters they belong to (as shown in Fig. 6A).**

### Supplementary Figures

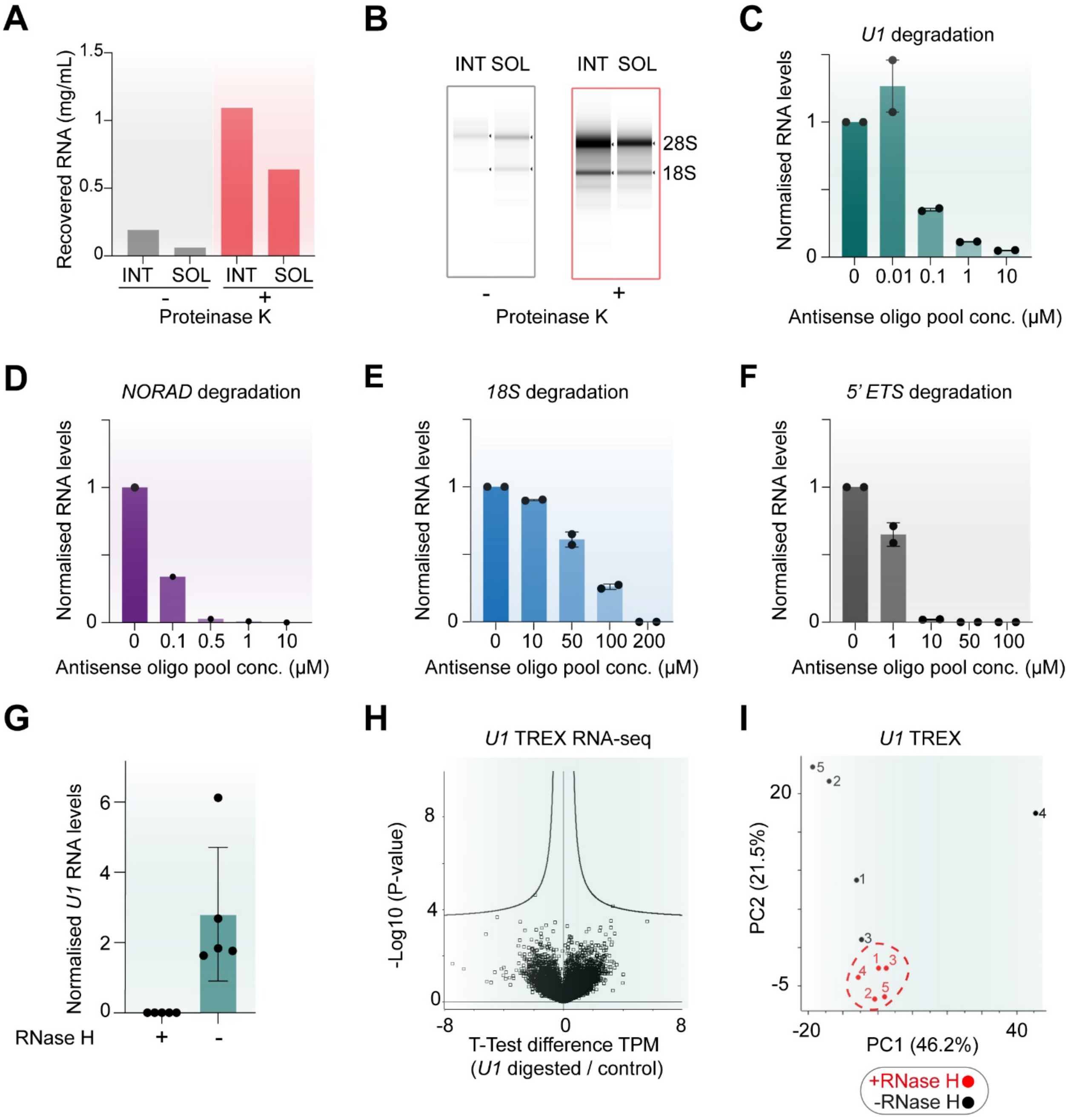


**Figure S1 (Related to Figure 1): Validation of TREX parameters.**

**(A)** Confirmation of the solubilization of intact RNA-protein adducts from the interface of crosslinked HCT116 cells. UV-C crosslinked HCT116 cells were lysed in TRIZOL and subjected to organic phase separation to isolate RNA-protein adducts in the interface (INT). The Interface pellets were subsequently solubilized according to the TREX protocol (SOL), before treatment with or without proteinase-K to remove the crosslinked proteins. The mixtures were then subjected to standard TRIZOL-based RNA extraction. Equal volumes from equal starting amounts for each INT or SOL sample, with or without proteinase K treatment, were quantified for RNA content using Nanodrop. Interface primarily contains protein-bound

RNA, since the majority of RNA is only recoverable after proteinase K digestion. More than half of this protein-bound RNA is solubilized by the TREX resolubilization protocol.

**(B)** Analysis of the integrity of the recovered RNA from (A) by capillary electrophoresis. Equal volumes from equal starting amounts for each INT or SOL sample, with or without proteinase K treatment, were resolved by capillary electrophoresis using Tapestation. The solubilized protein-bound RNAs are largely intact, as judged by visualization of full-length 28S and 18S rRNA bands.

**(C)** Assessment of the RNase H-mediated degradation of *U1* snRNA in solubilized interface fractions, by dose-dependent addition of tiling antisense DNA oligonucleotides. Solubilized protein-bound RNAs from the interface of phase separated UV-C-treated HCT116 cells were annealed to increasing concentrations of a pool of tiling DNA oligonucleotides complementary to the *U1* sequence. The annealed interface samples were then treated with RNase H to trigger the degradation of DNA-hybridized RNAs. The remaining amount of *U1* snRNA in each sample was then quantified by RT-qPCR, using specific RT-qPCR probes against *U1*. As input control, probes against *RPS18* and *GAPDH* were used, and the *U1* levels relative to the two controls were determined using the ΔCt method.

**(D)** Assessment of the RNase H-mediated degradation of *NORAD* lncRNA in solubilized interface fractions, by dose-dependent addition of tiling antisense DNA oligonucleotides. Solubilized protein-bound RNAs from the interface of phase separated UV-C-treated HCT116 cells were annealed to increasing concentrations of a pool of tiling DNA oligonucleotides complementary to the *NORAD* ND4 segment. The annealed interface samples were then treated with RNase H to trigger the degradation of DNA-hybridized RNAs. The remaining amount of *NORAD* ND4 segment in each sample was quantified by RT-qPCR, using specific PCR probes against this section of *NORAD*. As input control, probes against *RPS18* and *GAPDH* were used, and the *NORAD* ND4 levels relative to the two controls were determined using the ΔCt method.

**(E)** Assessment of the RNase H-mediated degradation of *18S* rRNA in solubilized interface fractions, by dose-dependent addition of tiling antisense DNA oligonucleotides. Solubilized protein-bound RNAs from the interface of phase separated UV-C-treated HCT116 cells were annealed to increasing concentrations of a pool of tiling DNA oligonucleotides complementary to the *18S* sequence. The annealed interface samples were then treated with RNase H to trigger the degradation of DNA-hybridized RNAs. The amount of *18S* in each sample was quantified by RT-qPCR, using specific PCR probes against *18S*. As input control, probes against *RPS18* and *GAPDH* were used, and *18S* levels relative to the two controls were determined using the ΔCt method.

**(F)** Assessment of the RNase H-mediated degradation of *5ETS* pre-rRNA region in solubilized interface fractions, by dose-dependent addition of tiling antisense DNA oligonucleotides. Solubilized protein-bound RNAs from the interface of phase separated UV-C-treated HCT116 cells were annealed to increasing concentrations of a pool of tiling DNA oligonucleotides complementary to the *5ETS* pre-rRNA sequence. The annealed interface samples were then treated with RNase H to trigger the degradation of DNA-hybridized RNAs. The amount of *5'ETS* pre-rRNA in each sample was quantified by RT-qPCR, using specific PCR probes against *5'ETS*. As input control, probes against *RPS18* and *GAPDH* were used, and *5'ETS* levels relative to the two controls were determined using the ΔCt method.

**(G)** Analysis of depletion efficiency in the RNase H-treated vs. untreated samples of the *U1* TREX experiment (related to Fig. 1C). Total RNA was extracted from aliquots of RNase H treated and untreated TREX samples , and was subjected to RT-qPCR analysis using specific PCR probes against *U1.* As input control, probes against *RPS18* and *GAPDH* were used, and *U1* levels relative to the two controls were determined in each sample using the ΔCt method. RNAse H treatment results in near complete *U1* removal.

**(H)** Analysis of depletion specificity in the RNase H-treated vs. untreated samples of the *U1* TREX experiment (related to Fig. 1C). Total extracted RNA from (G) was analyzed by whole-transcriptome RNA-seq to reveal differences in the transcriptome following RNase H treatment. TPM values for the identified RNAs were subjected to two-sample t-test analysis with permutation-based FDR calculation, to reveal differentially expressed RNAs between the RNase H treated and untreated samples. No transcripts were

significantly reduced by the RNase H treatment (note that U1 snRNA itself is not detectable in this type of RNA-seq due its small size). Curved lines mark the significance boundary (FDR = 0.05, S0 = 0.1).
**(I)** Principal component analysis (PCA) of the LFQ values from the proteomics analysis of RNase H treated (red) and untreated (black) *U1* TREX samples. Five biological replicates per condition were analyzed.

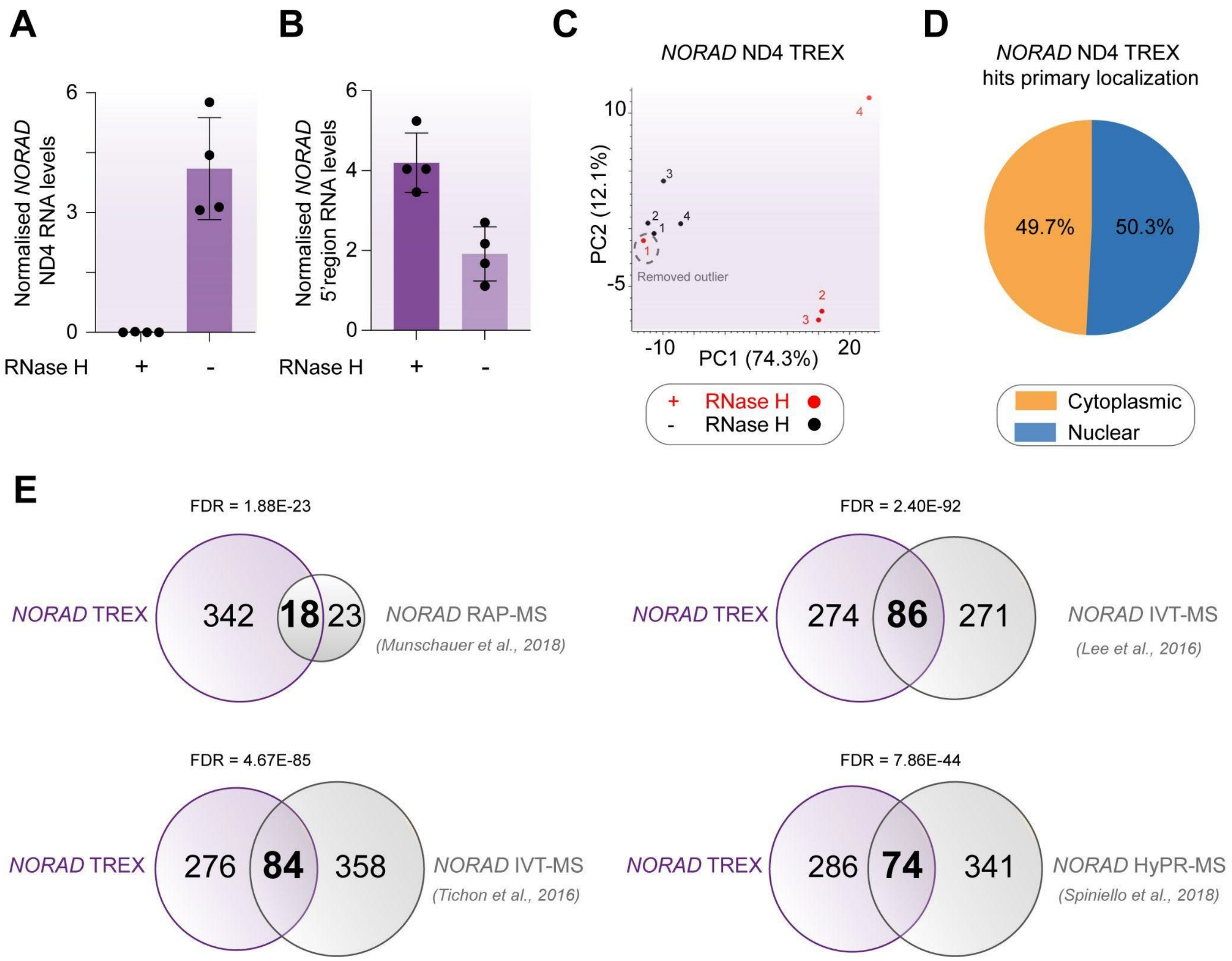


**Figure S2 (Related to Figure 2): Analysis of *NORAD* by TREX.**

**(A)** Analysis of depletion efficiency in the RNase H-treated vs. untreated samples of the *NORAD* ND4 TREX experiment (related to Fig. 2B). Total RNA was extracted from aliquots of RNase H treated and untreated TREX samples, and was subjected to RT-qPCR analysis using specific RT-qPCR probes against the *NORAD* ND4 domain*.* As input control, probes against *RPS18* and *GAPDH* were used, and *NORAD* ND4 levels relative to the two controls were determined in each sample using the ΔCt method. RNAse H treatment results in near complete degradation of *NORAD* ND4 segment.

**(B)** Analysis of the depletion region-specificity in the NORAD ND4 TREX samples (related to Fig. 2B). The total RNA extracted from *NORAD* ND4 TREX samples in (A) was subjected to RT-qPCR analysis using specific RT-qPCR probes against the 5'end segment of *NORAD.* As input control, probes against *RPS18* and *GAPDH* were used, and the levels *NORAD* 5' region relative to the two controls were determined in each sample using the ΔCt method. RNAse H treatment does not degrade the 5' end segment of *NORAD.*

**(C)** PCA of the LFQ values from the proteomics analysis of RNase H treated (red) and untreated (black) *NORAD* ND4 TREX samples. Four biological replicates per condition were analyzed. RNase H-treated sample 1 is an outlier and groups much more closely with the untreated samples, suggesting experiment failure.

**(D)** GOCC analysis of the primary subcellular location associated with the significant hits from *NORAD* ND4 TREX. While 49.7% of the hits were annotated as belonging to the 'nuclear part' category of GOCC, 50.3% were annotated as belonging to the 'cytoplasmic part' category.

**(E)** Venn diagram of the overlap between the lists of *NORAD* ND4 hits identified by TREX, and four previous *NORAD* interactome capture studies (Lee et al., 2016; Munschauer et al., 2018; Spiniello et al., 2018; Tichon et al., 2016). A highly significant overlap, calculated using Fisher's exact test with Benjamini-Hochberg FDR estimation, was detected in each comparison. The exact number of overlapping vs. non-overlapping proteins in each comparison, as well as the Fisher's exact test FDR values, are depicted on the Venn diagrams.

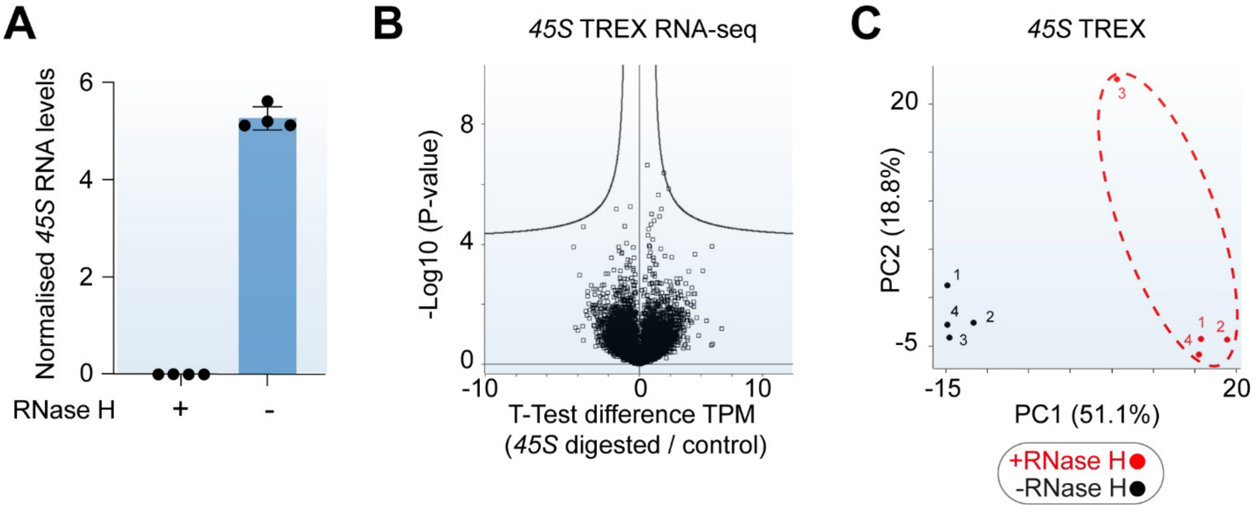


**Figure S3 (Related to Figure 3): Analysis of *45S* by TREX.**

**(A)** Analysis of depletion efficiency in the RNase H-treated vs. untreated samples of the *45S* rRNA TREX experiment (related to Fig. 3B). Total RNA was extracted from aliquots of RNase H treated and untreated TREX samples, and was subjected to RT-qPCR analysis using specific RT-qPCR probes against two segments of *45S rRNA.* As input control, probes against *RPS18* and *GAPDH* were used, and *45S* rRNA levels relative to the two controls were determined in each sample using the ΔCt method. RNAse H treatment results in near complete degradation of *45S* rRNA.

**(B)** Analysis of depletion specificity in the RNase H-treated vs. untreated samples of the *45S* TREX experiment (related to Fig. 3B). Total extracted RNA from (A) was analyzed by whole-transcriptome RNA-seq to reveal differences in the transcriptome following RNase H treatment. TPM values for the identified RNAs were subjected to two-sample t-test analysis with permutation-based FDR calculation, to reveal differentially expressed RNAs between the RNase H treated and untreated samples. No transcripts were significantly reduced by the treatment (note that rRNA itself is not detectable in this assay due to the ribo-depletion procedure used for library preparation). Curved lines mark the significance boundary (FDR = 0.05, S0 = 0.1).

**(C)** PCA of the LFQ values from the proteomics analysis of RNase H treated (red) and untreated (black) *45S* rRNA TREX samples. Four biological replicates per condition were analyzed.

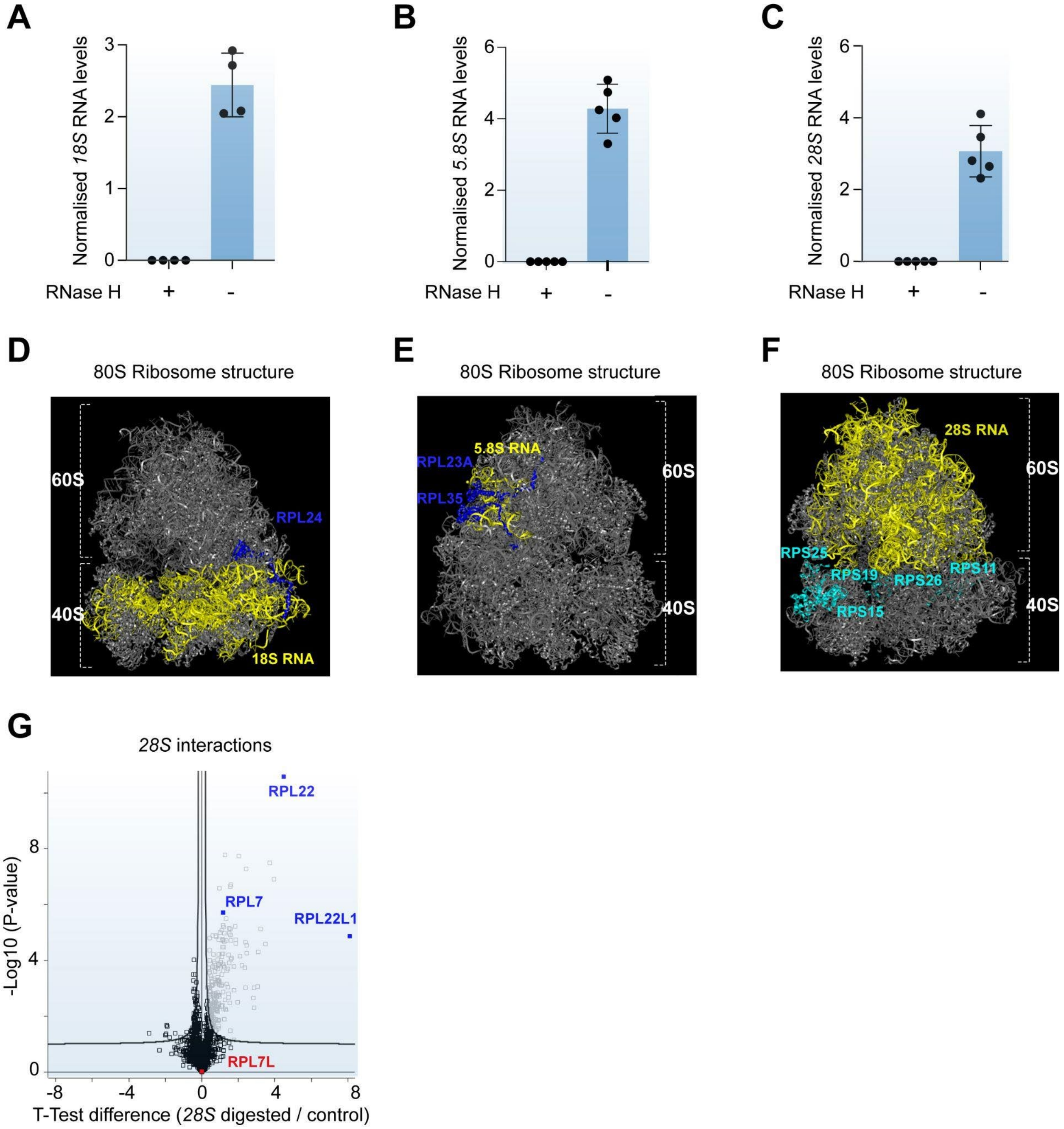


**Figure S4 (Related to Figure 4): Analysis of *18S, 5.8S,* and *28S* rRNA interactomes by TREX.**

**(A)** Analysis of depletion efficiency in the RNase H-treated vs. untreated samples of the *18S* rRNA TREX experiment (related to Fig. 4B). Total RNA was extracted from aliquots of RNase H treated and untreated TREX samples, and was subjected to RT-qPCR analysis using specific RT-qPCR probes against the *18S* segment of *45S* rRNA*.* As input control, probes against *RPS18* and *GAPDH* were used, and *18S* rRNA levels relative to the two controls were determined in each sample using the ΔCt method. RNAse H treatment results in near complete degradation of *18S* rRNA.

**(B)** Analysis of depletion efficiency in the RNase H-treated vs. untreated samples of the *5.8S* rRNA TREX experiment (related to Fig. 4D). Total RNA was extracted from aliquots of RNase H treated and untreated

TREX samples, and was subjected to RT-qPCR analysis using specific RT-qPCR probes against the *5.8S* segment of *45S* rRNA. As control, probes against the *18S* segment and *GAPDH* were used, and the relative *5.8S* rRNA levels were determined in each sample using the ΔCt method. RNAse H treatment results in near complete degradation of the *5.8S* segment.

**(C)** Analysis of depletion efficiency in the RNase H-treated vs. untreated samples of the *28S* rRNA TREX experiment (related to Fig. 4E). Total RNA was extracted from aliquots of RNase H treated and untreated TREX samples, and was subjected to RT-qPCR analysis using specific RT-qPCR probes against the *28S* segment of *45S* rRNA. As control, probes against the *18S* segment and *GAPDH* were used, and the relative *28S* rRNA levels were determined in each sample using the ΔCt method. RNAse H treatment results in near complete degradation of the *28S* segment.

**(D)** Structure of the Human 80S ribosome (*PDB ID: 4UG0*), visualized on PyMOL, with the *18S* rRNA (yellow) and RPL24 (blue) molecules highlighted on the structure. RPL24 extends from the 60S subunit into the 40S subunit and makes extensive contacts with *18S* rRNA.

**(E)** Structure of the Human 80S ribosome (*PDB ID: 4UG0*), visualized on PyMOL, with the *5.8S* rRNA (yellow), RPL23A (blue), and RPL35 (blue) molecules highlighted on the structure. Both RPL23A and RPL35 make extensive direct contacts with the *5.8S* rRNA in the 60S subunit.

**(F)** Structure of the Human 80S ribosome (*PDB ID: 4UG0*), visualized on PyMOL, with the *28S* rRNA (yellow), RPS11 (cyan), RPS15 (cyan), RPS19 (cyan), RPS25 (cyan), and RPS26 (cyan) highlighted on the structure. All highlighted proteins are located at the interface of the two ribosomal subunits.

**(G)** Volcano plot of the two-sample t-test comparison of *28S* rRNA digested vs undigested TREX samples, with RPL22, RPL22L1, RPL7, and RPL7L highlighted on the graph. Curved lines mark the significance boundary (FDR = 0.05, S0 = 0.1). Both RPL22 and RPL22L1 paralogues are detected amongst the significant *28S* rRNA interactors, while only RPL7 is detected as an interactor.

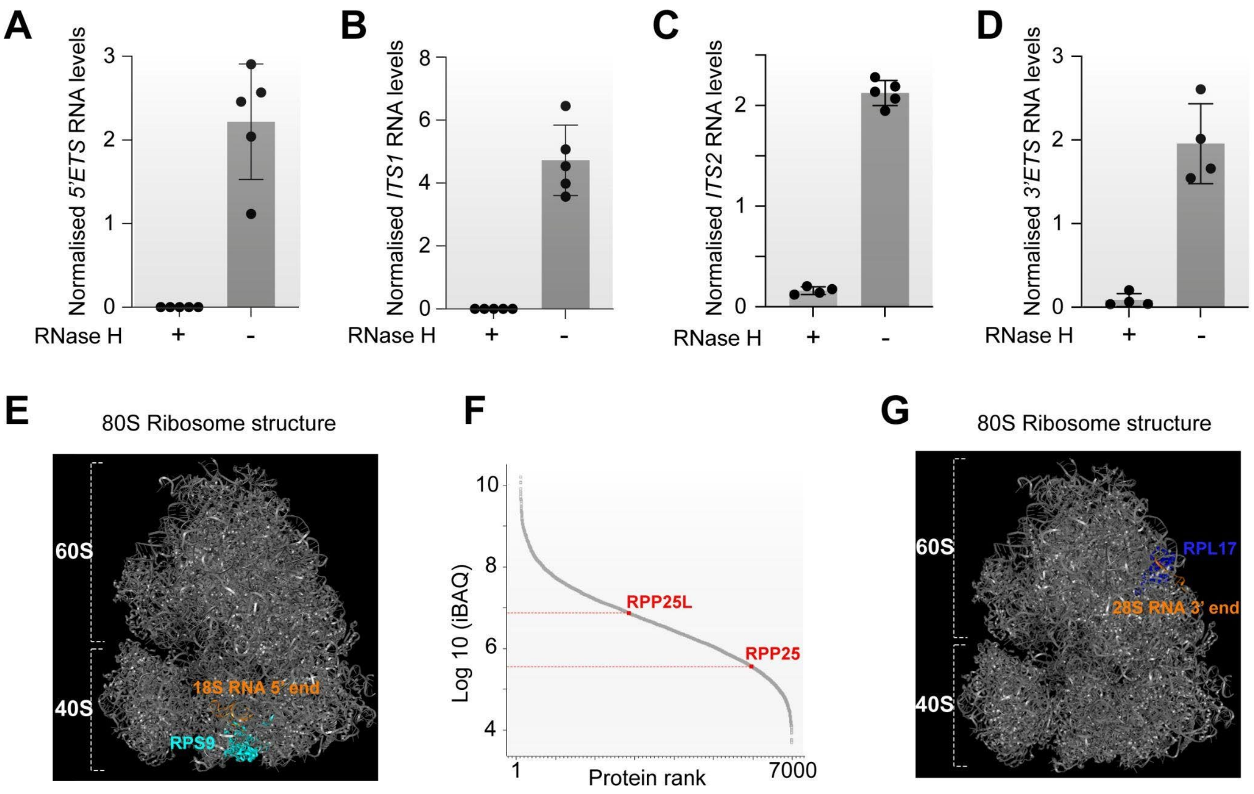


**Figure S5 (Related to Figure 5): Analysis of 5'*ETS, ITS1, ITS2, and 3'ETS* interactomes by TREX.**

**(A)** Analysis of depletion efficiency in the RNase H-treated vs. untreated samples of the *5'ETS* rRNA TREX experiment (related to Fig. 5B). Total RNA was extracted from aliquots of RNase H treated and untreated TREX samples, and was subjected to RT-qPCR analysis using specific RT-qPCR probes against the *5'ETS* segment of *45S* rRNA*.* As control, probes against the *18S* segment and *GAPDH* were used, and the relative *5'ETS* rRNA levels were determined in each sample using the ΔCt method. RNAse H treatment results in near complete degradation of the *5'ETS* segment.

**(B)** Analysis of depletion efficiency in the RNase H-treated vs. untreated samples of the *ITS1* rRNA TREX experiment (related to Fig. 5D). Total RNA was extracted from aliquots of RNase H treated and untreated TREX samples, and was subjected to RT-qPCR analysis using specific RT-qPCR probes against the *ITS1* segment of *45S* rRNA*.* As control, probes against the *18S* segment and *GAPDH* were used, and the relative *ITS1* rRNA levels were determined in each sample using the ΔCt method. RNAse H treatment results in near complete degradation of the *ITS1* segment.

**(C)** Analysis of depletion efficiency in the RNase H-treated vs. untreated samples of the *ITS2* rRNA TREX experiment (related to Fig. 5F). Total RNA was extracted from aliquots of RNase H treated and untreated TREX samples, and was subjected to RT-qPCR analysis using specific RT-qPCR probes against the *ITS2* segment of *45S* rRNA*.* As control, probes against the *18S* segment and *GAPDH* were used, and the relative *ITS2* rRNA levels were determined in each sample using the ΔCt method. RNAse H treatment results in near complete degradation of the *ITS2* segment.

**(D)** Analysis of depletion efficiency in the RNase H-treated vs. untreated samples of the *3'ETS* rRNA TREX experiment (related to Fig. 5H). Total RNA was extracted from aliquots of RNase H treated and untreated TREX samples, and was subjected to RT-qPCR analysis using specific RT-qPCR probes against the *3'ETS* segment of *45S* rRNA*.* As control, probes against the *18S* segment and *GAPDH* were used, and the relative *3'ETS* rRNA levels were determined in each sample using the ΔCt method. RNAse H treatment results in near complete degradation of the *3'ETS* segment.

**(E)** Structure of the Human 80S ribosome (*PDB ID: 4UG0*), visualized on PyMOL, with the 5' end of *18S* rRNA (orange) and RPS9 (cyan) highlighted on the structure. RPS9 binds specifically at the 5'end of *18S*.
**(F)** The ranked plot of the iBAQ absolute protein abundance measurements from the total proteome of HCT116, revealing RPP25L to be > 20 fold more expressed than RPP25.
**(G)** Structure of the Human 80S ribosome (*PDB ID: 4UG0*), visualized on PyMOL, with the 3' end of *28S* rRNA (orange) and RPL17 (blue) highlighted on the structure. RPL17 binds specifically at the 3'end of *28S*.
